# Engineered Serum Markers for Noninvasive Monitoring of Gene Expression in the Brain

**DOI:** 10.1101/2022.07.17.500352

**Authors:** Sangsin Lee, Shirin Nouraein, James J. Kwon, Zhimin Huang, Jerzy O. Szablowski

## Abstract

Noninvasive efforts to map brain gene expression have been hampered by low sensitivity and limited access to the brain. Here, we introduce a new platform that enables multiplexed, noninvasive, and site-specific monitoring of brain gene expression through a novel class of engineered reporters called Released Markers of Activity (RMAs). Instead of detecting gene expression in the less accessible brain, RMA reporters exit from a known brain region into the blood, where they can be easily measured with biochemical techniques. Expressing RMAs at a single brain site, typically covering ∼1% of the brain volume, provides up to a 39,000-fold signal increase over the baseline *in vivo*. Further, expression of RMAs in as few as several hundred neurons was sufficient for their reliable detection. When placed under a promoter upregulated by neuronal activity, RMAs could be used to measure neuronal activity in specific brain regions with a simple blood draw. We found that chemogenetic activation of cells expressing *Fos*-responsive RMA increased serum levels of RMA over 4-fold compared to non-activated controls. By contrast, a control RMA expressed under a constitutive neuronal promoter did not show such upregulation, demonstrating multiplexed ratiometric measurement with RMAs and proving specificity of neuronal activity discrimination. Together, our study pioneers a new noninvasive paradigm for repeatable and multiplexed monitoring of gene expression in an intact brain with sensitivity that is currently unavailable through other noninvasive gene expression reporter systems.

## INTRODUCTION

Monitoring gene expression dynamics in the living brain is critical for studying its cellular activity, understanding complex cognitive behaviors^1-3^, and controlling the onset of neurological diseases^4,5^. However, the delicate and intricate architecture of the brain poses significant challenges to tracking gene expression *in vivo* and stresses the need for noninvasive analytical methods that are both sensitive and specific.

Several prominent noninvasive technologies have exploited the use of genetically encodable reporters for mapping gene expression in the intact brain. Although they vary in their operational complexity, each methodology must contend with trade-offs in terms of noninvasiveness, sensitivity, specificity, and depth. For example, magnetic resonance imaging (MRI) can visualize gene expression throughout the whole brain with sub-millimeter spatial resolution using genetically encoded contrast agents, including iron-binding^6-8^ and non-metallic reporters^9-11^. However, reporter sensitivity and contrast resolution are marred by competing background signals from surrounding tissue^12^. Additionally, efforts to measure endogenous promoter activity, including that of the immediate early gene (IEG) *Fos*, have yet to yield success. In ultrasound imaging, great strides have been made in enhancing contrast through the application of genetically encoded air-filled gas vesicles (GVs)^13,14^, which have also been shown to augment signal contrast in MRI^15^. Still, the use of GVs has several drawbacks, including having to deliver ultrasound to each GV variant, which is difficult in thick skulls, large transgene sizes, and limited multiplexing capabilities. Lastly, optical imaging systems, such as optoacoustic^16,17^ and fluorescence^18^ imaging utilize reporters to enable brain imaging with subcellular spatial resolution at millisecond timescales and have been widely adopted for *in vivo* interrogations of neuronal activity. However, the heterogeneity of the brain as well as the opacity of the skull inherently limit the depth at which optical modalities can probe gene expression activities. Because of the strong light scattering and absorption properties of brain tissue, optical systems face the formidable challenge of imaging subcortical and deep cortical regions, particularly in large brains^19^. While recent improvements in BLI have expanded detection into the deep brain, substrates still need to penetrate the blood brain barrier (BBB). In addition, the degree of multiplexity for BLI^20^ and fluorescent imaging *in vivo* has not yet reached the levels achieved by biochemical detection methods, such as antibody-^21^, DNA hybridization^22^, or mass spectrometry^23^-based assays.

Unsurprisingly then, most brain-wide gene expression studies are performed on post-mortem brains^24-26^, which involve tissue-destructive biochemical techniques (e.g. high-throughput RNA sequencing^27^) that ultimately preclude longitudinal assessments of the same animal. To date, no technology has been developed for monitoring brain gene expression that is (1) noninvasive, (2) sensitive enough to measure expression in a small number of cells anywhere in the brain, (3) repeatable in the same animal, (4) capable of simultaneously imaging multiple molecule types, and (5) inexpensive and accessible to many research laboratories.

Here, we describe a new class of gene expression reporters, called Released Markers of Activity (**RMAs**), for noninvasive, sensitive, site-specific, and repeatable measurement of gene expression in the intact brain. RMAs leverage two naturally occurring phenomena – first, secretion from the neuron to release RMAs into the interstitial space and, second, reverse transcytosis to allow RMAs to cross the BBB into the bloodstream^28^. Because RMAs enter the blood, they can be detected using any sensitive biochemical method. Moreover, RMAs are compatible with multiplexed biochemical detection assays, some of which can reach single molecule sensitivity^29-32^. Unlike other methodologies that measure the concentration of reporters within the brain, RMAs present a radical new approach to monitoring brain gene expression that can be achieved with a simple blood draw.

In this study, we expressed RMAs in multiple brain regions, including the striatum, hippocampus, and midbrain, and easily detected the reporters after a single viral injection. RMAs were measured at high levels in the plasma and were measurable even in the sub-thousand neuron range. Further, chemogenetic activation of specific brain regions led to an increase in RMA signals without significant change in constitutive gene expression, demonstrating that RMAs can be used to discriminate neuronal activity *in vivo*. Our work establishes RMAs as a novel paradigm for noninvasive monitoring of gene expression dynamics in the brain.

## RESULTS

### Engineering the RMA reporter as a serum marker for gene expression in the brain

The RMA platform involves genetically labelling targeted brain sites with synthetic BBB-permeable RMA reporters, allowing for their release into the blood, and subsequently measuring the level of plasma RMAs to quantify gene expression in the brain (**Fig. 1a**). An RMA that can be transported from the brain to the blood would need to cross two biological barriers: the neuronal cell membrane and the BBB. Therefore, we designed an RMA reporter that contains two functional protein domains – one to facilitate crossing the cell membrane and another for crossing the BBB.

**Figure 1.**
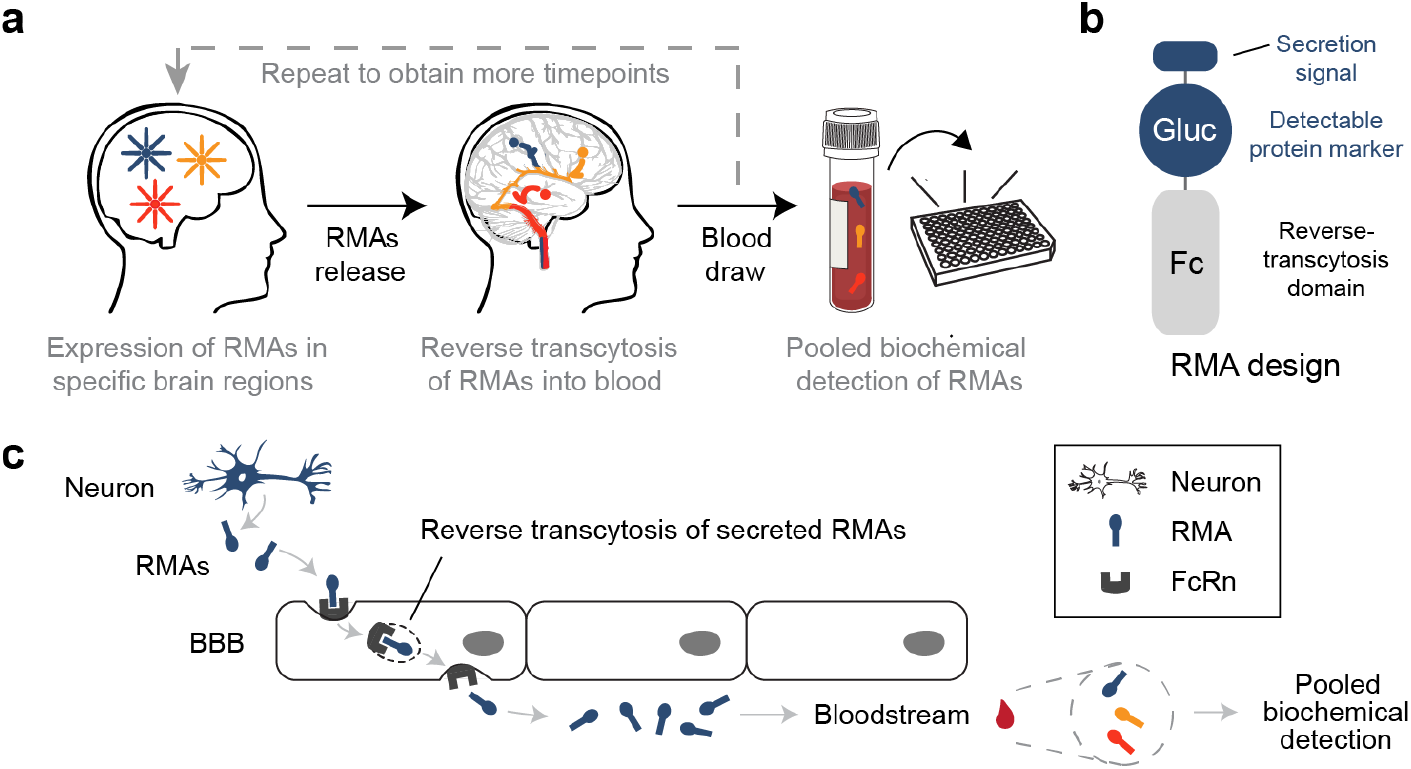
Noninvasive monitoring of gene expression in the brain with Released Markers of Activity (RMAs). (**a**) RMAs are a new class of genetically-encoded reporters of gene expression. RMAs are secreted from a known brain region into the interstitial space of the brain and then transported into the blood. Once in blood, they can be detected with any biochemical serum analysis method, without the confounds associated with imaging within solid tissues. The spatial information is tied to the molecular identity of an RMA and is provided by a known site of delivery of the RMA-encoding genes. **(b)** RMAs are proteins that contain a cell secretion signaling sequence, an easily detectable marker (e.g. luciferase, fluorescent protein, or an epitope of an antibody), and an Fc-region of an antibody that enables reverse transcytosis across the BBB. **(c)** An RMA (blue) is expressed in transduced cells (blue neuron) and secreted into the surrounding tissue. RMAs are fused to a moiety that recognizes neonatal Fc receptor (FcRn), which mediates the transport of RMAs into the blood. The process of transport is called reverse transcytosis. Once in blood, all RMAs in the sample can be detected using any biochemical method.

First, RMAs need to undergo exocytosis to move from the neuron into the extracellular space. To enable crossing the cell membrane, we chose to incorporate into the RMA design naturally secreted *Gaussia* luciferase (Gluc), a highly sensitive reporter that can also be used with BLI techniques,^33,34^ endowing the RMA with both a cell secretion sequence and a detectable protein marker domain (**Fig. 1b**).

Second, to enable RMAs to traverse the BBB, we leveraged an important feature of the reverse transcytosis that mediates the efflux of antibodies from the central nervous system (CNS) back into systemic circulation^35-37^. In this mechanism, the fragment crystallizable (Fc) region of an antibody binds to the neonatal Fc receptor (FcRn) expressed on the BBB in a pH-dependent manner. Fc binds avidly to FcRn at the endosomal pH (<6.5) in the CNS space but not at the physiological pH (7.4) in the blood^28,38^. The strict pH-dependent binding of Fc to FcRn thus, in effect, favors the unidirectional release of antibodies from the brain into the blood (**Fig. 1c**). To functionalize RMAs to undergo reverse transcytosis, we first selected Fc regions of three different immunoglobulin G (IgG) antibodies: the human IgG1 monomeric Fc (mFc) and mouse IgG1 and IgG2a Fcs. We then fused each Fc to the Gluc reporter to construct Gluc-Fc RMA variants.

### Secretion of RMAs by PC-12 cells

To assess RMA secretion from cells, we expressed the Gluc-Fc variants under the neuron-specific hSyn promoter in PC-12, a widely-used murine cell line for studying neurosecretion^39^. Subsequently, we measured the amount of secreted Gluc-Fc in the culture media by luciferase assay (**Fig. 2a and Supplementary Fig. 1a**). For each variant, we also tested a truncation mutant (Gluc-Fc Δa.a.1-17), which lacks the N-terminal secretion signal peptide. Our results showed that all Gluc-Fc variants with the signal peptide accumulated in the media over time, indicating that fusing Gluc to Fc does not compromise its ability to be secreted (**Fig. 2b and Supplementary Fig. 1b**). In addition, we observed that replacing the secretion signal peptide of native Gluc with that of the murine antibody (Igκ) preserves secretory function, suggesting that the secretion signal peptide is an interchangeable but essential component of efficient RMA release from the cells (**Supplementary Fig. 1b**).

**Figure 2.**
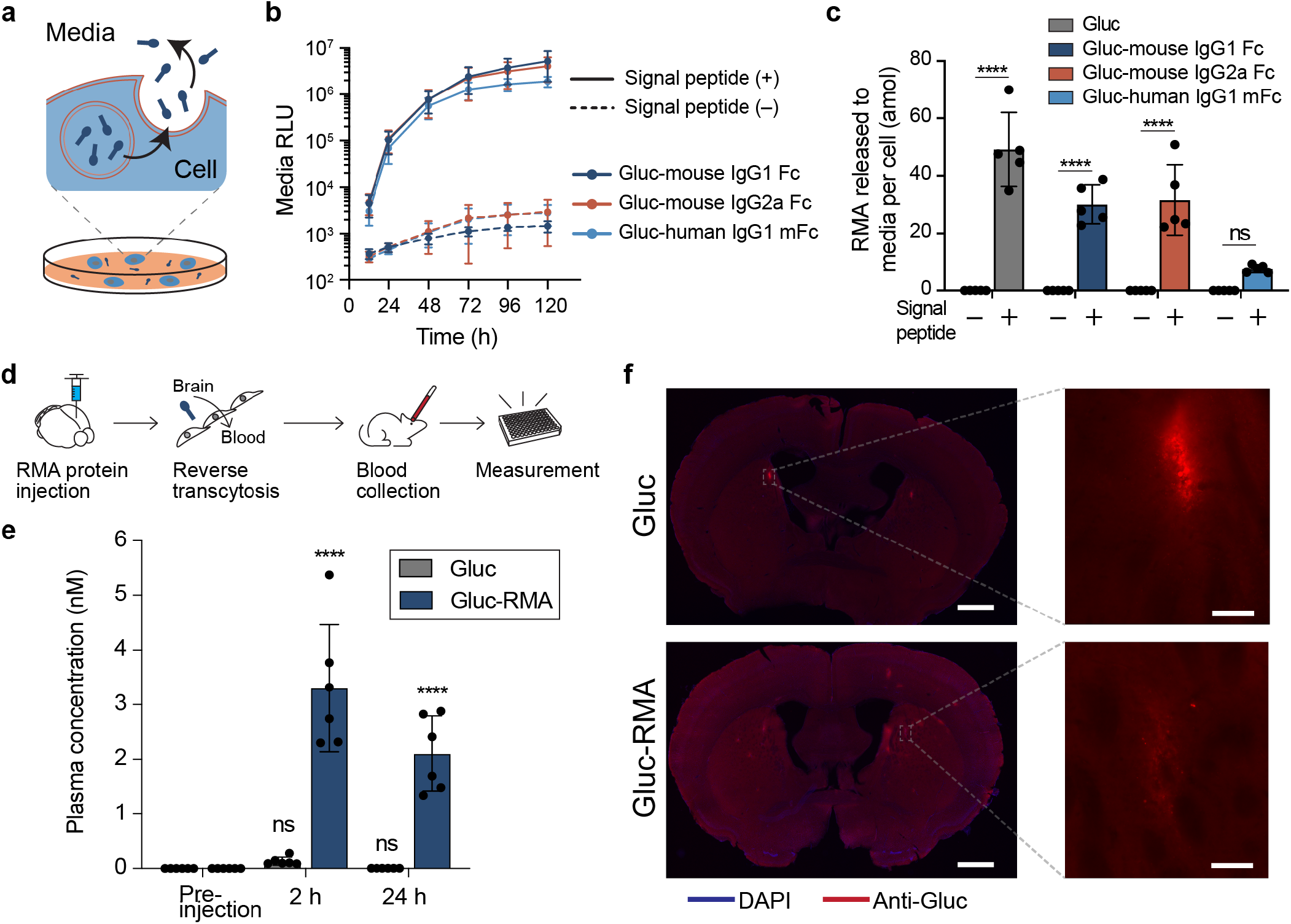
RMA reporters translocate from the brain into the bloodstream. **(a)** Schematic of RMA secretion *in vitro* for crossing the first barrier of the cellular membrane. **(b)** RMA secretion by PC-12 cells. Relative luminescence unit (RLU) values measured from the culture media reveal the signal peptide-dependent secretion of Gluc-Fc RMA variants. *n*=5 independent cultures analyzed. **(c)** Estimated amount of RMA proteins released per PC-12 cell after 72 hr post-transfection. \*\*\*\**P*<0.0001, ns (not significant) in comparison of each variant with and without the signal peptide, using Two-way ANOVA, Sidak’s test. **(d)** Experimental scheme for testing reverse transcytosis of RMA for crossing the BBB into the bloodstream. **(e)** Plasma concentration of RMAs measured from the collected blood after bilaterally injecting 1μL of 20 μM (20 pmol) RMA proteins per injection into CP of the mice brains. *n*=6 independent mice analyzed. \*\*\*\**P*<0.0001, ns (not significant) in comparison with the plasma concentration of the respective RMA measured from the pre-injection time point, using two-way ANOVA Sidak’s test. **(f)** Representative images of the stained mice brain slices showing the remaining RMA proteins in the brain after 24 hr post-injection. Scale, 1000 μm and 50 μm of the left (whole brain) and right (enlarged) images, respectively. All data are shown as mean ± SD.

Given the attomolar detection sensitivity of Gluc^40^, we then studied the secretion rates of the Gluc-Fc variants to estimate the minimum number of cells required for detection using luciferase assay. We bicistronically-expressed Gluc-Fc and GFP under the hSyn promoter in PC-12 cells and quantified the amount of Gluc-Fc released into the media. We then normalized the amount of Gluc-Fc per cell using the total number of GFP-positive cells (**Supplementary Fig. 1c**). On average, Gluc showed the highest secretion rate with 49.2 ±

12.8 amol per cell, followed by Gluc-mouse IgG2a Fc (31.6 ± 12.2), Gluc-mouse IgG1 Fc (30.1 ± 6.8), and Gluc-human IgG1 mFc (7.6 ± 1.2) (**Fig. 2c**). These secretion rates suggest that one PC-12 cell could be sufficient for readout and demonstrate detectability of our secreted RMAs *in vitro*. As expected, RMAs lacking the signal peptide showed no measurable secretion.

### RMAs exit from the brain into the blood

To examine whether RMAs can cross the BBB, we performed direct injections of RMA proteins into the caudate putamen (CP) of the mouse brain, where IgG efflux is mediated by the interaction between Fc and FcRn^35^. Among the RMA variants, we selected Gluc-mouse IgG1 Fc, hereafter referred to as Gluc-RMA, as it contains the Fc of the native host. We injected 20 pmol of either Gluc or Gluc-RMA into each hemisphere and measured the released reporters in blood samples (**Fig. 2d**). Within 2 h after the injection, we observed a significant rise in the plasma concentration of Gluc-RMA (24.4-fold higher than that of Gluc), suggesting that RMA transportation from the brain to the blood occurs through an Fc-dependent mechanism (**Fig. 2e-f and Supplementary Fig. 2**). Importantly, between 2 and 24 h, we observed that Gluc-RMA plasma levels decreased only by 35 ± 13%, whereas Gluc displayed a 96 ± 2% reduction close to the baseline (**Fig. 2e**). We also found the *β*-phase half-life (*t*_1/2_) of Gluc-RMA was substantially greater than that of Gluc (6,269 ± 2,283 min vs 31 ± 3 min) (**Supplementary Fig. 3**). These data are consistent with the pharmacokinetics of Fc-fusion proteins, wherein the fusion protein acquires extended *t*_1/2_ through its interaction with FcRn, which helps to reduce protein degradation^28,41^. Together, our data suggest that Gluc-RMA is released from the brain and circulates with multi-hour-long *t*_1/2_, allowing for its accumulation in blood.

### RMAs detect gene expression in as few as hundreds of neurons

After establishing that Gluc-RMA can traverse out of the brain and into the blood, we studied whether it could be used to detect brain gene expression *in vivo*. We injected adeno-associated virus (AAV) encoding both Gluc-RMA and GFP controlled under the constitutive neuronal hSyn promoter into the mouse brain and assayed the plasma for the released reporter (**Fig. 3a**). We first injected into both hemispheres of the CP and found respective 36,867- and 49,530-fold signal increases at 2 and 3 weeks post-delivery when compared with 0 weeks (baseline) (**Supplementary Fig 4a-b**).

Given these high Gluc-RMA signals, we next sought to examine possible regional dependencies of Gluc-RMA by singly injecting the CP, CA1, and substantia nigra regions located in the striatum, hippocampus, and midbrain, respectively (**Fig. 3b-d**). We found >20,000-fold higher signals over baseline in all three regions, demonstrating that gene expression in various local brain regions can lead to detectable RMA signal levels in the serum (**Fig 3e-g**), regardless of their location. Furthermore, the signal levels persisted up to the 3rd week, possibly due to plasma Gluc-RMA reaching steady-state, wherein the rate of production matches the rate of degradation under the constitutive expression.

To determine the fewest number of neurons that can be transduced and later discerned in histological images, we performed another injection into a single CP site using 1/1000^th^ of the initial AAV dose (see Methods). We observed a 43-fold signal increase over the baseline in approximately 815 (± 327, n=3) neurons as estimated using histological analysis (**Supplementary Fig. 4c-d**), suggesting that Gluc-RMA reliably detects ∼0.001% of neurons in the mouse brain^42^. To test the correlation between plasma signals and the number of transduced neurons or AAV dose, we collected and analyzed the data of those injected in the CP with different AAV doses. We discovered a linear relationship (r^2^=0.90) for up to 56,841 transduced neurons (∼0.1% of mouse neurons) (**Supplementary Fig. 4e-f**). Taken together, these results indicate that Gluc-RMA provides a linear dynamic range throughout all tested conditions with a sensitivity to monitor gene expression in as few as hundreds of neurons.

### High-sensitivity monitoring of brain cell type-specific gene expression using Cre lines

To observe gene expression in a selected brain cell type, we delivered hSyn-controlled double floxed Gluc-RMA-IRES-GFP into TH-Cre mice, which express Cre in dopaminergic neurons under the control of a tyrosine hydroxylase (TH) promoter. We specifically targeted the left ventral tegmental area (VTA), which is rich with TH-positive cells (**Fig. 4a**). At 2 and 3 weeks post-delivery, plasma Gluc-RMA signals were 1,022- and 1,409-fold higher than at 0 weeks, respectively (**Fig. 4b**). Expressions of Gluc-RMA and GFP were specific to TH-positive cells at the local injection site (**Fig. 4c**), providing evidence that Gluc-RMA can detect brain gene expression in small neuronal cell populations, and in a cell type-specific manner.

**Figure 3.**
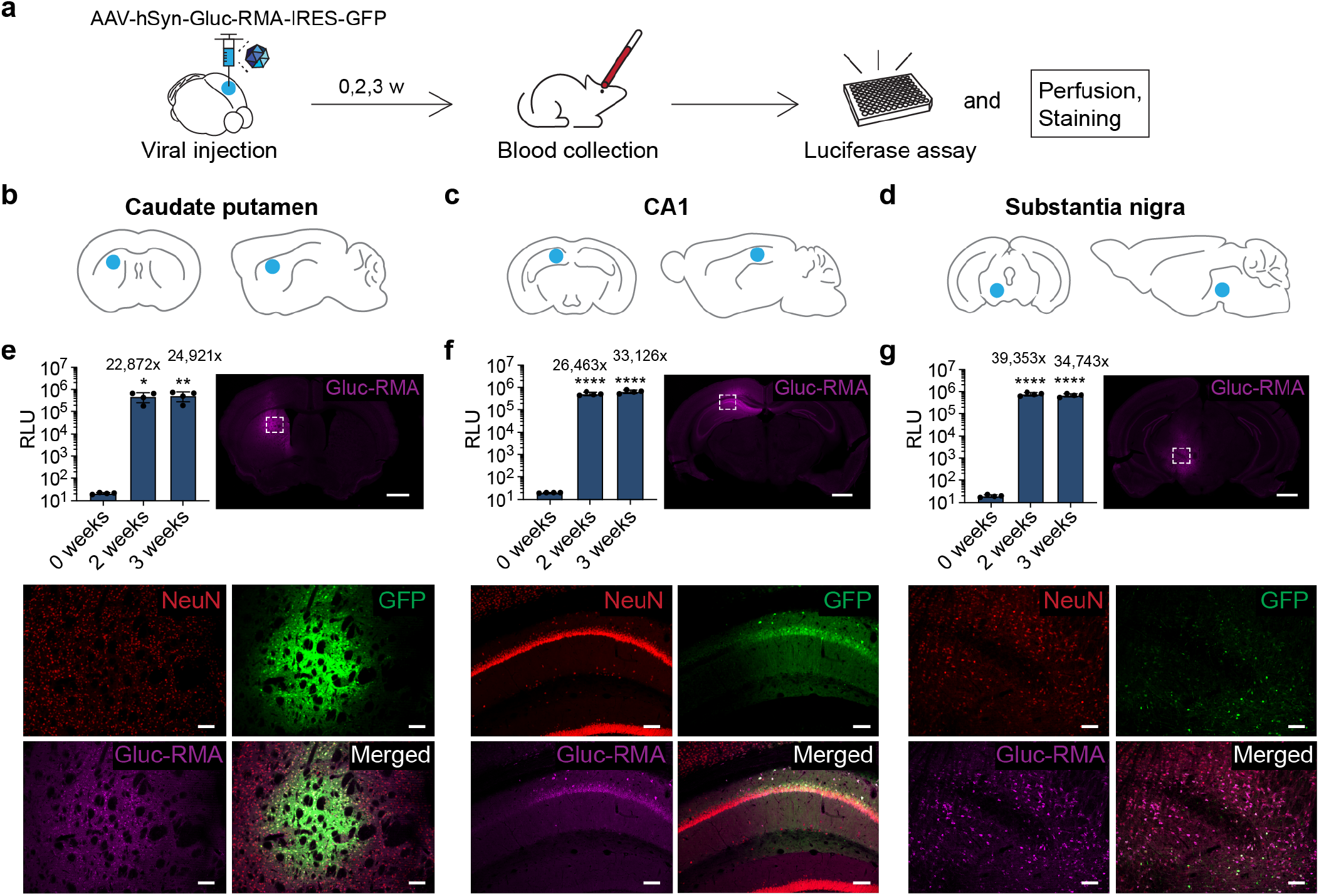
Gluc-RMA detects gene expression at multiple local regions. **(a)** Experimental scheme for detecting gene expression in brain regions. Mouse is injected with AAV encoding Gluc-RMA and subjected to blood collection for measurement of the released Gluc-RMA reporters. **(b), (c), and (d)** AAV injection sites. Left and right brain schemes show the coronal and sagittal views, respectively. Blue circles indicate the target injection sites: **(b)** caudate putamen (CP, striatum). **(c)** CA1 (hippocampus). **(d)** substantia nigra (SN, midbrain). **(e), (f)**, and **(g)** Plasma bioluminescence signal and representative images showing gene expression of Gluc-RMA in the target brain region -CP, CA1, and SN, respectively. Left (bar graph): RLU measured from the collected blood containing Gluc-RMA after injecting 2.4 × 10^9^ vg AAVs into the respective brain sites indicated in **(b), (c)**, and **(d)**. The number shown above each bar refers to the signal fold increase compared with the 0 weeks baseline. *n*=4 independent mice analyzed. \**P*<0.05, \*\**P*<0.01, \*\*\*\**P*<0.0001, in comparison with the signal at 0 weeks, using One-way ANOVA, Tukey’s test. Data are shown as mean ± SD. Right (whole-brain image): Gluc-RMA expression at the local injected sites. Scale, 1000 μm. Bottom (large images): Enlarged views of the white rectangular region indicated on the whole-brain image. Scale, 100 μm.

**Figure 4.**
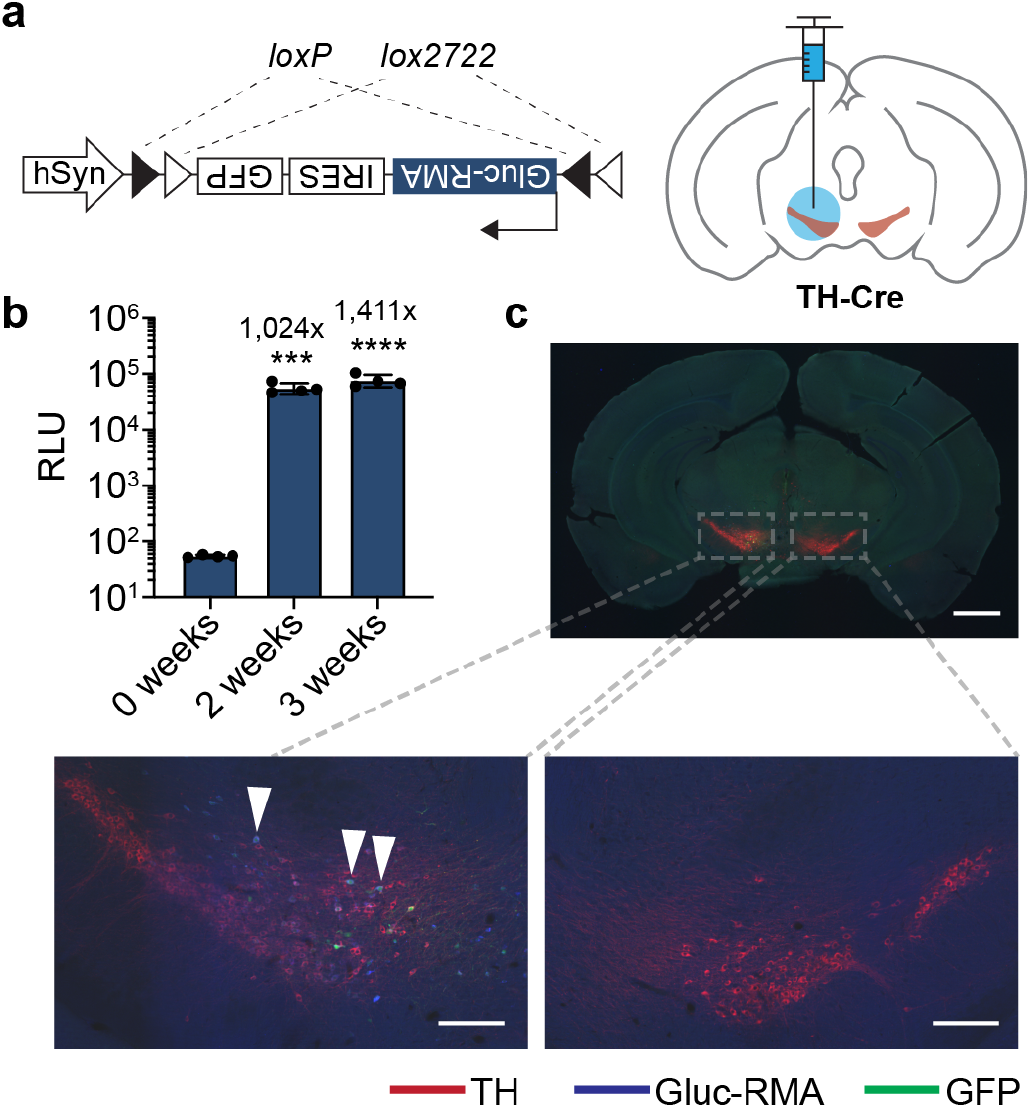
Detecting gene expression of specific brain cell types with high sensitivity using Gluc-RMA. (**a**) Schematic of selective expression of Gluc-RMA and GFP in the brain cells that express Cre recombinase. Cre recognizes the double-floxed gene (Gluc-RMA-IRES-GFP) flanked by the two *loxP* sites and inverts the sequence back to the correct orientation, which results in the expression of Gluc-RMA and GFP. Coronal view of the AAV injection site (blue) at the VTA region of the brain that expresses Cre (brown) in TH-Cre mice. **(b)** Plasma bioluminescence signal measured from the collected blood samples containing the released Gluc-RMA after injection of AAVs at the dose of 1.2 × 10^9^ vg. The number shown above each bar refers to the signal fold increase compared with the 0-week baseline. *n*=4 independent mice analyzed. \*\*\**P*<0.001, \*\*\*\**P*<0.0001, in comparison with the signal at 0 weeks, using One-way ANOVA, Tukey’s test. Data are shown as mean ± SD. **(c)** Representative image of stained brain slice showing expression of Gluc-RMA and GFP among TH positive brain cells. Several of the Gluc-RMA cells are highlighted with arrowheads. Scale, 1000 μm (whole-brain image), 200 μm (enlarged images).

### RMAs capture Fos gene expression activity

We next investigated the ability of RMAs to track changes in the expression of the IEG *Fos*, which rapidly expresses upon cellular stimulus or neuronal activity^2^. For this *in vitro* experiment, we transfected PC-12 with plasmid encoding Gluc-RMA controlled under the *Fos* promoter (Fos-Gluc-RMA) (**Supplementary Fig. 5a**). To induce *Fos* expression in transfected PC-12, we supplemented the culture media with nerve growth factor (NGF)^43^. Subsequently, we performed luciferase assays on the culture media from multiple time points. Upon NGF induction, we observed increased expression of Gluc-RMA and *Fos* (**Supplementary Fig. 5b-c**). Consistent with these results, the luminescence signal of the culture media rose significantly within 6 hr of exposure to NGF, suggesting that Gluc-RMA can generate a distinguishable signal output in response to changes in promoter activity (**Supplementary Fig. 5d**).

### Noninvasive measurement of neuronal activity in specific brain regions

To determine whether RMAs could be used to detect neuronal activity *in vivo*, we designed a double-conditional strategy to link Gluc-RMA expression to neuronal activity in the brain (**Fig.5a**). To enable chemogenetic neuromodulation, we implemented a DREADD (designer receptor exclusively activated by designer drug) system. For our purposes, we chose the excitatory DREADD hM3Dq^44^, which, when activated by intraperitoneally (i.p.)-administered clozapine-*N*-oxide (CNO), elicits robust neuronal firing^44,45^ and *Fos* accumulation^46^. For the readout of plasma Gluc-RMA, we incorporated a doxycycline (Dox)-dependent Tet-Off system called Robust Activity Marking (RAM)^47^ to couple the RMA reporter gene to a synthetic *Fos* promoter and gain temporal control over its transcription (**Fig. 5b**). Upon neuronal stimulation with CNO, both an active *Fos* promoter and the absence of Dox is needed to drive the expression of Gluc-RMA. To account for any variations in gene expression activities between mice, we also constructed an internal control *Cypridina* luciferase (Cluc)-RMA constitutively expressed under the hSyn promoter, thus demonstrating multiplexed monitoring capability of RMAs.

**Figure 5.**
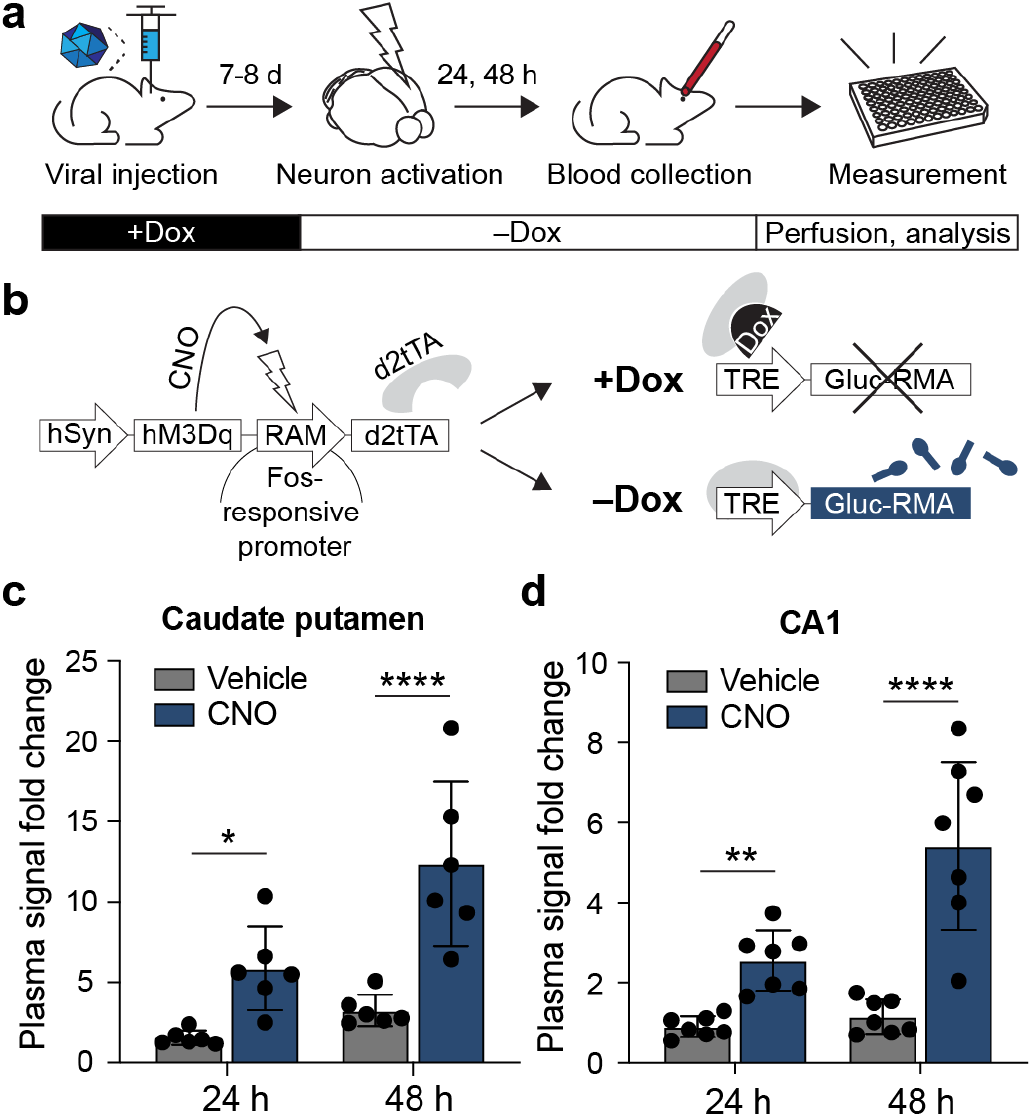
Gluc-RMA detects neuronal activity *in vivo*. (**a**) Experimental scheme for detecting neuronal activity through blood tests with RMAs. Mice undergo injection of AAVs carrying RMA reporter genes, then induction of neuronal activation, and blood collection for measurement of the released RMA reporter. **(b)** Schematic of the RAM system for detecting neuronal activity. Upon CNO injection, activatory hM3Dq DREADD induces *Fos* expression. Subsequently, the RAM promoter drives the expression of d2tTA transactivator. If Dox is absent the d2tTA then binds to TRE and induces the expression of Gluc-RMA. If present, Dox prevents d2tTA from binding to TRE and thus prevents the expression, allowing for temporally-gated recording of neuronal activity. **(c)** Plasma bioluminescence signal fold change of Gluc-RMA after inducing chemogenetic activation at CP of the striatum. Signals were normalized by dividing the RLU values obtained from Gluc-RMA over Cluc-RMA. **(d)** Plasma signal after inducing chemogenetic activation at CA1 of the hippocampus. *n*=6 (for CP) and *n*=7 (for CA1) independent mice analyzed, using two-way ANOVA, Sidak’s test. \**P*<0.05, \*\**P*<0.01, \*\*\**P*<0.001, \*\*\*\**P*<0.0001. Data are shown as mean ± SD. RAM: robust activity marking, d2tTA: tetracycline-controlled transactivator fused to a degradation domain, TRE: tTA-responsive element, Dox: doxycycline.

We then prepared AAVs encoding the *Fos*-responsive, RAM-controlled Gluc-RMA and delivered them into the left CP of mice, along with Cluc-RMA and hM3Dq (**Supplementary Fig. 6a**). We fed mice a Dox chow diet and replaced it with a Dox-free diet 48 hr prior to administering CNO for neuronal activation (**Fig. 5a**). Our results showed that mice injected with CNO revealed plasma Gluc-RMA signals that were 3.8-fold higher than the vehicle at 48 hr post activation, without significant change in the expression of Cluc-RMA (**Supplementary Fig. 6b-d**). We observed a similar trend in the CA1 of the hippocampus, with a 4.6-fold increase in the plasma signal (**Fig. 5e and Supplementary Fig. 7**). Collectively, our findings demonstrate the multiplexed ratiometric measurements of RMAs as well as their ability to discriminately report on *in vivo* neuronal activity of specific brain regions.

### RMA enhances in vivo bioluminescence imaging (BLI)

We next explored whether Gluc-RMA could be used to improve BLI. *In vivo* imaging system (IVIS) allows for noninvasive imaging of cells or tissues of interest using optical sensors. Due to the poor penetration of light *in vivo*, researchers commonly rely on albino or nude animals^34^ or high concentrations of reporters^48^ and typically limit their studies to small animal species. Additionally, the choice of luminophore is limited, by the need of those molecules to cross the BBB. Because Gluc-RMA can be released from the brain and has a long *t*_1/2_ in the blood, we hypothesized that Gluc-RMA could be used with BLI to facilitate the measurement of gene expression levels within the brain areas transduced with Gluc-RMA.

We expressed Gluc or Gluc-RMA in the whole brain by intravenously (i.v.) injecting BBB-permeable PHP.eB AAV^49^ and then imaged mice or analyzed their plasma using IVIS or luciferase assay, respectively (**Fig. 6a and Supplementary Fig. 8a**). Gluc-RMA improved the IVIS photon emission by a factor of 102 over Gluc, which showed detectable but not significant signal against the wild-type (**Fig. 6b**). A similar trend was observed in the plasma assays showing 193-fold improvement over Gluc (**Fig. 6c**). Furthermore, we confirmed that Gluc-RMA signals are correlated to gene expression levels through our AAV dose response analysis (**Fig. 6d and Supplementary Fig. 8b**). Taken together, these data suggest that Gluc-RMA improves the imaging performance of BLI by substantially enhancing its signal intensity.

**Figure 6.**
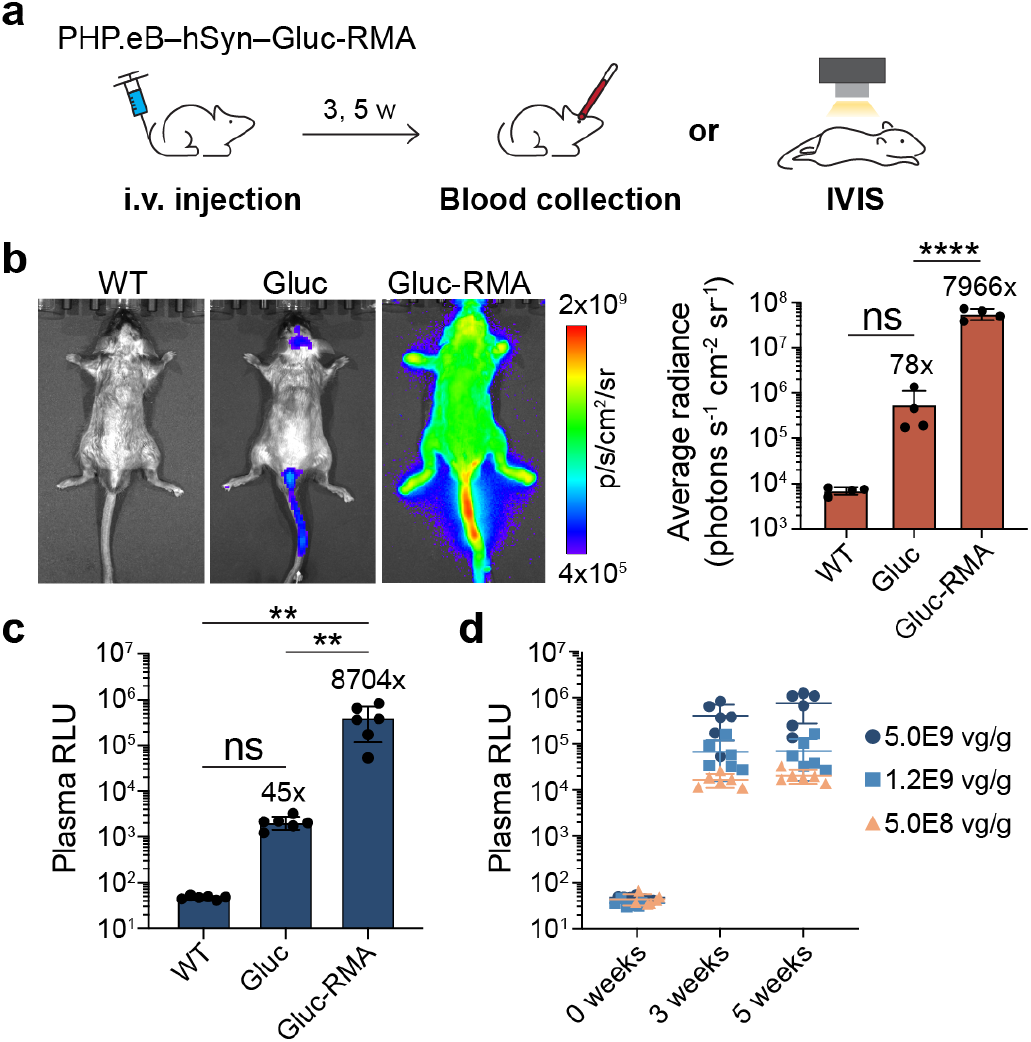
Gluc-RMA enhances bioluminescence imaging. (**a**) Schematic of delivery of the BBB-permeable PHP.eB virus encoding Gluc-RMA to the brain through intravenous (i.v.) injection and subsequent collection of the blood or BLI for measurement of the brain gene expression. **(b)** Bioluminescence imaging of mice taken after 3 weeks post-PHP.eB delivery at the dose of 5.0 × 10^9^ vg g^-1^ mouse. 12 μmol kg^-1^ of native coelenterazine was injected through the tail-vein and immediately imaged under the IVIS. Average radiance of the upper body was quantified using the Living Image software. *n*=4 independent mice analyzed. **(c)** Plasma RLU measured at 3-week post-PHP.eB delivery of the same viral dose. **(d)** PHP.eB dose response to the plasma bioluminescence signal. For **(b)** and **(c)**, \*\**P*<0.01, \*\*\*\**P*<0.0001, ns (not significant), using one-way ANOVA, Tukey’s multiple comparison test. For **(c)** and **(d)**, *n*=6 independent mice analyzed. Data are shown as mean ± SD. WT: wild type.

## DISCUSSION

Here, we establish RMAs as a new class of reporters to noninvasively measure gene expression in the brain. The current version of RMAs is optimized for high-sensitivity detection. To achieve this, we repurposed an endogenous pathway that allowed us to transport RMAs from the brain to the blood. The presence of the Fc region in the reporter provides a dual function of enabling the RMA to cross the BBB and prolonging its lifetime in the blood. RMAs accumulate in the blood over time and, owing to their long half-life, avoid the rapid clearance or low concentrations commonly encountered by natural brain-derived biomarkers^50^. RMAs are thus versatile gene expression reporters that can be expressed under any promoter of interest to monitor long-term changes in gene expression or efficiency of gene delivery to the brain in individual animals. Such long-term gene expression changes can be observed, for example, in tracking the dynamics of neuronal subtypes^51^, brain disorder pathogenesis^52,53^, aging^54,55^, or transgene expression following gene therapy administration^56^.

Since the brain is completely vascularized, RMAs can be used in any region of the brain, regardless of whether that region is deep or cortical. Thus, RMAs avoid the obstacles faced by many other methodologies that are limited by the depth of penetration, tissue scattering, or skull absorption of the penetrant waves used to image reporters. We have demonstrated signal levels between ∼20,000-40,000-fold over the baseline in three commonly studied brain regions after a single intracranial injection of AAVs carrying RMAs. RMAs could be of utility in large animal models where tissue scattering or skull absorption preclude the use of optical systems, such as intravital BLI. Because of its physical accessibility, blood has been commonly used for diagnosing various medical conditions, including cancer^34^ and neurodegenerative diseases^50,57-60^. Therefore, if effective in humans, blood assays for RMA could be clinically adopted to provide patients with a more convenient option over noninvasive techniques that image directly on the brain. Lastly, RMAs democratize access to noninvasive measurement of gene expression in the brain, opening this technique for use, for example, in high-throughput screening scenarios. Readout of RMAs does not require complicated scanners, such as MRI, because it relies on serum chemistry that is accessible to many research laboratories.

We demonstrated multiplexed RMA readout using Gluc-RMA and Cluc-RMA, which react with different substrates to emit different bioluminescence signals. We selected Gluc or Cluc for their ability to be secreted and conveniently assayed, but one could instead construct a compound of secreted library proteins fused to Fc to implement a highly multiplexed RMA system. In combination with highly multiplexed protein detection method, such as mass spectrometry^61^, RMAs have the potential to achieve higher multiplexity than currently available noninvasive methodologies. If such multiplexed monitoring is implemented, RMAs could be used to independently monitor large numbers of cells or genes. This ability for high multiplexity could confer an advantage on using RMAs over other techniques that rely on the limited number of reporter variants or fluorescent channels available.

The ability to monitor gene expression activities of individual or a subset of neuronal populations with high sensitivity will help with discriminating different functional networks in the deep brain. Our histological assessment suggested that expression of RMAs in 815 neurons was sufficient to produce 43-fold higher RMA signals compared with the baseline (**Supplementary Fig. 4d**). We note that the experiment with the 1,000-fold reduction in AAV dose could have resulted in a lower MOI (multiplicity of infection) relative to our high dose experiments, suggesting that sensitivity could be greater in some cases, e.g. transduction of sparse cell populations with high doses of the AAV. The plasma signals (around 2×10^3^ RLU) were one order of magnitude lower than the expected RLU values (815 neurons × 26.0 ≈ 2×10^4^ RLU) based on its linear relationship to the number of transduced neurons (**Supplementary Fig. 4e**), indicating that gene expression levels per neuron were comparatively low with the reduced AAV dose. These results support the idea that Gluc-RMA could report on even fewer neurons if small numbers of cells can be transduced with a high concentration of the AAV.

The difference in the levels of plasma Gluc-RMA after chemogenetic activation began to appear after 24 h, consistent with the time taken for fluorescent reporters to express in the brain cells using the RAM system^47,62^. These data suggest that the expression of the reporter protein, including transcription and translation processes, accounts for the major time delay in detecting the output signals. In addition, given their long half-life, the current RMAs are less applicable to transient neuronal activities or studies that require high temporal resolution and more useful for measuring persistent changes in the timescale of days. To increase the temporal resolution, the important work ahead would be to accelerate the expression of RMAs and shorten their lifespan in the blood without compromising the Fc-dependent release from the brain. However, such modification would likely reduce the sensitivity of the readout. The application of emerging single-molecule protein detection methods^63^ could allow RMAs to both achieve faster readout kinetics and maintain the high sensitivity observed in this study.

With the RMA platform, we have pioneered the use of blood as an alternative route for measuring gene expression in the intact brain. RMAs are genetically-encodable reporters that exhibit high sensitivity, repeatability, and multiplexity, making them well-suited for numerous neuroscience applications, such as monitoring differential gene expression activities among cell type-, circuit-, or spatial-specific brain cells, observing long-term changes in different neuronal subtypes, or monitoring changes in neuronal activity. We anticipate that by opening a new noninvasive pathway into the brain, RMAs will prove to be an invaluable and promising tool for brain gene expression studies.

## MATERIALS AND METHODS

### Animal subjects

Wild-type C57BL/6J (Strain #000664) and transgenic TH-Cre (Strain #008601) male and female mice at 8-10 weeks old were purchased from the Jackson Laboratory. Animals were housed with a 12 h light-dark cycle and were provided with food and water ad libitum. All animal experiments were performed under the protocol approved by the Institutional Animal Care and Use Committee of Rice University.

### Plasmid construction

To construct AAV-hSyn-RMA (Gluc-Fc), the vector AAV-hSyn-GlucM23-iChloC-EYFP (Addgene #114102) was digested with KpnI and EcoRV (New England Biolabs) to isolate the backbone containing the hSyn promoter. GlucM23, a Gluc variant, was amplified by PCR from the same vector and its DNA was extracted using the Monarch DNA Gel Extraction Kit (New England Biolabs). DNA segments for Fc regions, including the human IgG1 Fc (Addgene #145165), and mouse IgG1 Fc (Addgene #28216) and IgG2a Fc (Addgene #114492), were amplified and extracted similarly. Gluc alone or Gluc with Fc was inserted into the digested backbone through Gibson Assembly. To make mFc from the human IgG1 Fc, the relevant mutations were introduced using site-directed mutagenesis^64^. For RMA controls that lack the signal peptides, the first 17 amino acids of the Gluc sequence were skipped during the amplification. To co-express RMA and GFP, IRES-GFP sequence from the bicistronic vector (Addgene #105533) was amplified and inserted downstream of the RMA coding region to construct AAV-hSyn-RMA-IRES-GFP. Plasmid for RMA controlled under the Fos promoter AAV-Fos-RMA was constructed by extracting the Fos promoter from the plasmid Fos-tTA (Addgene #34856) and replacing the hSyn with the Fos promoter from our AAV-hSyn-RMA. For plasmids that encode Cluc-RMA (Cluc-Fc), the Cluc DNA was obtained from pClucIPZ (Addgene #53222) and used instead of Gluc for assembly.

To make pET-T7-RMA-His for purifying RMA proteins, the RMA sequence was amplified from our AAV-hSyn-RMA. His tag was attached to the C-terminus of RMA using reverse primers containing the overhang that encodes six His residues. The pET28a vector was kindly provided by the Tabor Lab at Rice University. The amplified DNA was then inserted into the pET28a backbone using Gibson Assembly.

To construct AAV-hSyn-hM3Dq-RAM-d2tTA, AAV-hSyn-hM3Dq-mCherry (Addgene #50474) was digested with SalI and PmlI to obtain the backbone that contains the hSyn promoter. hM3Dq was amplified separately to add an HA tag to its N-terminus. RAM-d2tTA was amplified and extracted from AAV-RAM-d2tTA-TRE-MCS (Addgene #63931). Two inserts HA-hM3Dq and RAM-d2tTA were then assembled into the backbone. For AAV-TRE-RMA-IRES-GFP, AAV RAM-d2tTA-TRE-MCS was digested with NheI and KpnI and the segment RMA-IRES-GFP was used as an insert for Gibson Assembly.

### Cell culture for luciferase assay

PC-12 (ATCC) was cultured in RPMI 1640 medium (Corning) supplemented with heat inactivated 10% horse serum (Life Technologies) and 5% fetal bovine serum (FBS) (Corning). Cells were incubated in humidified air with 5% CO2 at 37 ^°^C and split every 2 d with a subcultivation ratio of 1:2 or 1:3.

For *in vitro* luciferase assay, PC-12 was seeded at 200,000 cells per well in a 12-well plate. After 16-20 h, 1,500 ng of plasmids encoding hSyn-RMA and 3.0 μl of lipofectamine 2000 (Life Technologies) were used to transfect PC-12 following the manufacturer’s protocol. Then, 25 μl of the culture media were collected at different time points and stored in –20 ^°^C until use. For Gluc substrate, 0.5 mM native coelenterazine (CTZ) stock (Nanolight Technology) was dissolved in N_2_-purged acidified DMSO (0.06 N HCl) and stored at –80 ^°^C. Before measuring bioluminescence, the CTZ stock was diluted to 20 μM in luciferase assay buffer (10 mM Tris, 1 mM EDTA, 1.2 M NaCl, pH 8.0) and kept in dark at room temperature for 1 h. Media samples were thawed in ice and transferred to black 96-well plate (Corning). Infinite M Plex microplate reader (Tecan) was used to inject 50 μl of the assay buffer containing CTZ and measure the photon emission integrated over 30 s. Values were averaged to obtain the light unit per second.

### Measurement of RMA release per cell

PC-12 was transfected using the plasmid hSyn-RMA-IRES-GFP. After 72 h, the total cell number in each well was estimated using 10 μl of the culture media mixed with Trypan Blue and by counting the viable cells under the hemocytometer (INCYTO). The remaining cells and media were then centrifuged and processed separately. Cells were analyzed by flow cytometer (Sony SA3800) with at least 50,000 events to obtain the percentage of GFP positive cells. Non-transfected PC-12 was used to gate the positive cells (**Supplementary Fig. 9**) The number of transfected cells was calculated by multiplying the total cell number to the GFP^+^ percentage. For processing the media, luciferase assay was conducted to obtain their bioluminescence signals and a standard curve was generated using the fresh culture media spiked with purified RMA proteins at different concentrations. The total secreted RMAs was calculated by multiplying the concentration obtained from the standard curve to the total media volume. Finally, the number of RMAs released per cell was calculated by dividing the secreted RMAs over the number of transfected cells.

### Protein purification

Shuffle T7 Express chemically competent *E. coli* cells (New England Biolabs) were transformed using the plasmid pET-T7-RMA-His. The next day, a single colony was transferred into 3 ml of the LB medium as a starter culture to grow overnight at 30 ^°^C under 250 RPM in a shaker. The culture was then transferred into 1 L Terrific broth and grown in the same condition until the optical density at 600 nm reached 0.5. The culture flask was cooled on ice for 30 min, supplemented with 100 μM IPTG, and induced for 20 h at 16 ^°^C under 180 RPM. Cells were harvested by centrifugation at 4,000*g* for 20 min, resuspended in lysis buffer (300 mM NaCl, 50 mM NaH_2_PO_4_, 10 mM imidazole, 10% glycerol, pH 8.0) supplemented with ProBlock Gold protease inhibitor (Gold Biotechnology), and lysed by the sonicator (VCX 130, Sonics and Materials). Lysates were centrifuged at 12,000*g* for 30 min at 4 ^°^C. The resulting supernatant was subject to binding with Ni-NTA agarose resin (Qiagen) for 30 min at 4 ^°^C with gentle rotation and loaded into the glass chromatography columns (Bio-Rad) for wash and elution through gravity flow. Protein-bound resin was washed sequentially using lysis buffer with incremental increase of imidazole concentrations. Bradford protein assay (ThermoFisher Scientific) was performed throughout the washing procedure to check for the presence of non-specific proteins. RMA proteins were eluted using lysis buffer containing 500 mM imidazole and buffer exchanged into PBS using the Amicon centrifugal filter unit with 10 kDa cutoff (MilliporeSigma). The final protein in PBS was analyzed by the SDS-PAGE. BCA protein assay (ThermoFisher Scientific) was used to determine protein concentration.

### Adeno-associated virus production

PHP.eB AAV virus was packaged by adopting a previously published protocol with slight modifications^65^. In brief, HEK293T cells (ATCC) were transfected with the transfer plasmid PHP.eB iCap (Addgene #103005), and pHelper plasmids. After 24 h, cells were exchanged with a fresh DMEM (Corning) supplemented with 5% FBS and non-essential amino acids (Life Technologies). At 4 d post-transfection, cells were harvested, and media were mixed with 1/5 volume of PEG solution (40% PEG 8,000, 2.5 M NaCl) to precipitate AAV at 4 ^°^C for 2 h. Cells were resuspended in PBS and lysed by the freeze-thaw method. The precipitated AAV was pelleted by centrifugation, resuspended in PBS, and combined with the lysed cells. The combined lysate was added with 50 U ml^-1^ of Benzonase (Sigma-Aldrich) and incubated at 37 ^°^C for 45 min before being stored at –20 ^°^C for no more than one week.

AAV purification was carried out by the iodixanol gradient ultracentrifugation. Quick-seal tube (Beckman Coulter) was loaded with the iodixanol gradients (Sigma-Aldrich), including 60%,40%, 25%, and 15%. The frozen lysate was thawed and centrifuged at 2,000*g* for 10 min. The resulting clarified lysate was transferred on top of the iodixanol layers drop-by-drop. The tube was sealed and centrifuged at 58,400 RPM for 2.5 h using the 70 Ti fixed-angle rotor of an ultracentrifuge (Beckman Coulter). AAV was collected by extracting the 40%-60% iodixanol interface and washed using the Amicon centrifugal filter unit with 100 kDa cutoff (MilliporeSigma). The final AAV was filtered by passing through the 0.22 μm PES membrane. Viral titers were determined using the qPCR method.

### Stereotaxic injection

Protein or AAV was injected into the mice brains using a microliter syringe equipped with a 34-gauge beveled needle (Hamilton) installed to a motorized pump (World Precision Instruments) using a stereotaxic frame (Kopf). To inject RMA protein, 20 μM of RMA was injected bilaterally to CP (AP +0.25 mm, ML ±2.0 mm, DV –3.2 mm, 1 μl per hemisphere) infused at a rate of 200 nl min^-1^ and the needle was kept in place for 5 min before taking it out from the injection site. For AAV injections, PHP.eB serotype was used for all experiments. To deliver AAV encoding hSyn-RMA-IRES-GFP, 2.4 × 10^9^ vg in 200 nl was injected per site at 600 nl min^-1^ to the following coordinates: CP in the striatum (AP +0.25 mm, ML +2.0 mm, DV –3.2 mm), CA1 in the hippocampus (AP –1.94 mm, ML +1.0 mm, DV –1.3 mm), and substantia nigra in the midbrain (AP –3.28 mm, ML +1.5 mm, DV –4.3 mm). For conducting the 1/1000^th^ dilution of the initial dose (2.4 × 10^6^ vg), 50 nl was injected at 150 nl min^-1^ into the same CP coordinate. For testing the TH-Cre mice, 1.2 × 10^9^ vg of AAV encoding hSyn-DIO-RMA-IRES-GFP was injected into the left VTA (AP – 2.9 mm, ML +0.8 mm, DV –4.55 mm).

For chemogenetic neuromodulation experiments, mice were placed on 40 mg kg^-1^ of Dox chow (Bio-Serv) 24 h prior to surgery. AAV doses used are as follows: 2.0 × 10^9^ vg (hSyn-hM3Dq-RAM-d2tTA), 2.0 × 10^9^ vg (TRE-Gluc-RMA-IRES-GFP), and 1.1 × 10^9^ vg (hSyn-Cluc-RMA). For each surgery, AAV cocktail was prepared in 450 nl and injected over 1 min. The needle was kept at the injection site for 10 min owing to the relatively high volume of the cocktail. Dox chow was removed 48 h prior to inducing chemogenetic activation.

### Blood collection for luciferase assay

Mice were anesthetized in 1.5%-2% isoflurane in air or O_2_. Following, 1-2 drops of 0.5% ophthalmic proparacaine were applied topically to the cornea of an eye. Heparin-coated microhematocrit capillary tube (Fisher Scientific) was placed into the medial canthus of the eye and the retro-orbital plexus was punctured to withdraw 50-100 μl of blood. The collected blood was centrifuged at 1,500*g* for 5 min to isolate plasma and stored at –20 ^°^C until use. To conduct luciferase assay, 5 μl of plasma was mixed with 45 μl of PBS + 0.001% Tween-20 in a black 96-well plate. Using the microplate reader, bioluminescence of Gluc-RMA or Cluc-RMA was measured by injecting 50 μl of 20 μM CTZ or 1.0 μM vargulin (Nanolight Technology), respectively, dissolved in the luciferase assay buffer into the plasma sample.

### Half-life measurement

Mice were administered i.v. with 2 mg kg^-1^ of purified RMA proteins. Retro-orbital blood collections were performed at specific time points. Concentrations of RMAs in the collected blood were determined using the standard curves generated by the luciferase assays conducted on the normal mouse serum (Sigma-Aldrich) added with RMAs at known concentrations. To determine the pharmacokinetic parameters, AUC_inf_ (area under the curve to infinity) was calculated using the log-linear trapezoid method. Half-life (*t*_1/2_) of the distribution (*α*) or elimination (*β*) phase in the two-phase clearance model was calculated by applying the log-linear regression to the concentration data. The concentrations of the first three and the subsequent time points were used for determining the *α*- and *β*-phase *t*_1/2_, respectively. Clearance was calculated by dividing the dose by AUC_inf_.

### Drug administration

Water-soluble CNO (Hello Bio #HB6149) was dissolved in saline (Hospira) at 1mg ml^-1^ and stored at –20 ^°^C until use. To induce chemogenetic activation of mice expressing hM3Dq, CNO was injected i.p. at 5 mg kg^-1^, respectively. For the vehicle groups, saline was injected i.p. at the volume dose of 5 ml kg^-1^.

### Histological imaging and analysis

Mice brains were extracted and postfixed in 10% neutral buffered formalin (Sigma-Aldrich) overnight at 4 ^°^C. Coronal sections were cut at a thickness of 50 μm using a vibratome (Leica) and stored at 4 ^°^C in PBS. Sections were stained as follows: 1) block for 2 h at room temperature with blocking buffer (0.2% Triton X-100 and 10% normal donkey serum in PBS); 2) incubate with primary antibody overnight at 4 ^°^C; 3) wash in PBS for 15 min 3 times; and 4) incubate with secondary antibody for 4 h at room temperature. After last washes in PBS, sections were mounted on glass slides using the mounting medium (Vector Laboratories) with or without DAPI and cured overnight in dark at room temperature. Antibodies and dilutions used are as follows: rabbit anti-Gluc (1:1,500, Nanolight Technology), mouse IgG2a anti-Fos (1:500, Santa Cruz), mouse IgG2b anti-NeuN (1:1,500, Novus Biologicals), chicken IgY anti-TH (1:1,000, Aves Labs), mouse IgG1 anti-HA-594 (1:500, Life Technologies), and Alexa 350, 488, 594, 647 secondary antibodies (1:500, Life Technologies).

All images were acquired by the BZ-X810 fluorescence microscope (Keyence). Manual cell counting was performed using the Zen software (Zeiss) by a blinded observer uninformed of the experimental conditions. For the images obtained from the chemogenetic experiments, Gluc-RMA and GFP positive expression areas were quantified using ImageJ by stacking the TIFF image to RGB and recording the total area above the set brightness threshold (50 for Gluc-RMA and 34 for GFP).

### Estimation of the number of transduced cells

Every fifth of the 50 μm brain section (2 to 6 sections analyzed per brain) was taken to manually count the Gluc-RMA positive cells. For each section, cell density was estimated by dividing the number of counted cells over the transduction volume (the area occupying the transduced cells multiplied by the 50 μm thickness of the section). The total transduction volume in the brain was calculated using the Cavalieri method^66^, in which the average of the transduction volumes measured in all sections was multiplied by the rostro-caudal distance between the first and the last analyzed sections. Finally, the number of transduced cells was calculated by multiplying the average cell density by the total transduction volume.

### In vitro Fos activation

*Fos* activation in PC-12 was carried out by inducing with nerve growth factor (NGF). First, 18 mm-diameter glass coverslips were incubated in 10 μg ml^-1^ of collagen IV (Corning) in PBS and coated overnight at 4 ^°^C, then dried under UV light. The coated coverslips were placed onto a 12-well plate and PC-12 was seeded at 100,000 cells per well. After 20 h, 1500 ng of plasmids encoding AAV-Fos-RMA and 3 μl of lipofectamine 2000 were used to transfect PC-12. The next day, fresh media supplemented with 100 ng ml^-1^ of NGF was added to induce *Fos* activation. Then, 20 μl of media were collected at different timepoints and used for luciferase assay. After the last collection, cells on coverslips were fixed and stained against Gluc and Fos and imaged under the fluorescence microscope.

### IVIS spectrum imaging

Mice were anesthetized in 1.5%-2% isoflurane in air or O_2_. The body hair was removed using a hair trimmer. Mice were transferred to the IVIS spectrum imager (Perkin Elmer) while maintaining the appropriate anesthetic condition. For BLI settings, we used open filter, large binning, F/1 aperture control, and the auto-exposure time. We observed most images being generated with the 0.5 sec exposure time. Mice were injected i.v. with 12 μmol kg^-1^ of CTZ (Nanolight Technology, #303-INJ) and immediately imaged. Images were analyzed by the Living Image software (Caliper Life Sciences) to quantify the average radiance of the upper body and the head.

### Statistical analysis

Two-tailed *t* test with unequal variance was used to compare two data sets. One-way ANOVA with Tukey honestly significant difference post hoc test was used to compare means between more than two data sets. Two-way ANOVA with Sidak’s multiple comparison tests were used to compare data sets with two or more variables. Linear regression was used to find correlation between the plasma signal and the number of transduced neurons. All *P* values were determined using Prism (GraphPad Software), with the statistical significance represented as ns (not significant), \**P* < 0.05, \*\**P* < 0.01, \*\*\**P* < 0.001, \*\*\*\**P* < 0.0001.

## Data availability

The authors declare that all data supporting the results in this study are available within the paper and its Supplementary Information. The raw and analyzed datasets are available from the corresponding author upon reasonable request.

## ACKNOWLEDGEMENTS

The authors thank J.J. Tabor’s laboratory (Rice University) for providing the pET28a bacterial expression plasmid, and G. Bao’s laboratory (Rice University) for providing the usage of the ultracentrifuge. We thank V. Gradinaru laboratory (California Institute of Technology) and Caltech CLOVER Center for providing the pUCmini-iCAP-PHP.eB and pHelper plasmids. This research was supported by the David and Lucile Packard Foundation 2021-73005 (J.O.S.).

## AUTHOR CONTRIBUTIONS

J.O.S and S.L. conceived and planned the research. S.L. and J.O.S designed the experiments and wrote the manuscript with input from all other authors. S.L. performed and participated in all experiments described in the study. S.N. performed AAV production, stereotaxic injection, and retro-orbital blood collection. S.N. and J.J.K. performed the histological experiments. J.J.K. maintained PC-12, conducted the *in vitro Fos* activation, and analyzed histological images. S.N. and Z.H. assisted with IVIS imaging and drug administration. Z.H. constructed plasmids.

## REFERENCES

1 Richiardi, J. et al. Correlated gene expression supports synchronous activity in brain networks. Science 348, 1241–1244 (2015).

2 Minatohara, K., Akiyoshi, M. & Okuno, H. Role of immediate-early genes in synaptic plasticity and neuronal ensembles underlying the memory trace. Frontiers in molecular neuroscience 8, 78 (2016).

3 Pantazatos, S. P. et al. Whole-transcriptome brain expression and exon-usage profiling in major depression and suicide: evidence for altered glial, endothelial and ATPase activity. Molecular psychiatry 22, 760–773 (2017).

4 Wertz, M. H. et al. Genome-wide in vivo CNS screening identifies genes that modify CNS neuronal survival and mHTT toxicity. Neuron 106, 76-89.e78 (2020).

5 Seidlitz, J. et al. Transcriptomic and cellular decoding of regional brain vulnerability to neurogenetic disorders. Nature communications 11, 3358 (2020).

6 Genove, G., DeMarco, U., Xu, H., Goins, W. F. & Ahrens, E. T. A new transgene reporter for in vivo magnetic resonance imaging. Nature medicine 11, 450–454 (2005).

7 Shapiro, M. G. et al. Directed evolution of a magnetic resonance imaging contrast agent for noninvasive imaging of dopamine. Nature biotechnology 28, 264–270 (2010).

8 Sigmund, F. et al. Bacterial encapsulins as orthogonal compartments for mammalian cell engineering. Nature communications 9, 1990 (2018).

9 Desai, M., Slusarczyk, A. L., Chapin, A., Barch, M. & Jasanoff, A. Molecular imaging with engineered physiology. Nature communications 7, 13607 (2016).

10 Schilling, F. et al. MRI measurements of reporter-mediated increases in transmembrane water exchange enable detection of a gene reporter. Nature biotechnology 35, 75–80 (2017).

11 Mukherjee, A., Wu, D., Davis, H. C. & Shapiro, M. G. Non-invasive imaging using reporter genes altering cellular water permeability. Nature communications 7, 13891 (2016).

12 Farhadi, A., Sigmund, F., Westmeyer, G. G. & Shapiro, M. G. Genetically encodable materials for non-invasive biological imaging. Nature Materials 20, 585–592 (2021).

13 Farhadi, A., Ho, G. H., Sawyer, D. P., Bourdeau, R. W. & Shapiro, M. G. Ultrasound imaging of gene expression in mammalian cells. Science 365, 1469–1475 (2019).

14 Shapiro, M. G. et al. Biogenic gas nanostructures as ultrasonic molecular reporters. Nature nanotechnology 9, 311–316 (2014).

15 Lu, G. J. et al. Acoustically modulated magnetic resonance imaging of gas-filled protein nanostructures. Nature materials 17, 456–463 (2018).

16 Gottschalk, S. et al. Rapid volumetric optoacoustic imaging of neural dynamics across the mouse brain. Nature biomedical engineering 3, 392–401 (2019).

17 Lauri, A. et al. Whole-cell photoacoustic sensor based on pigment relocalization. ACS sensors 4, 603–612 (2019).

18 Marvin, J. S. et al. A genetically encoded fluorescent sensor for in vivo imaging of GABA. Nature methods 16, 763–770 (2019).

19 Ntziachristos, V. Going deeper than microscopy: the optical imaging frontier in biology. Nature methods 7, 603–614 (2010).

20 Iwano, S. et al. Single-cell bioluminescence imaging of deep tissue in freely moving animals. Science 359, 935–939 (2018).

21 de Wildt, R. M. T., Mundy, C. R., Gorick, B. D. & Tomlinson, I. M. Antibody arrays for high-throughput screening of antibody–antigen interactions. Nature biotechnology 18, 989–994 (2000).

22 Shendure, J. & Ji, H. Next-generation DNA sequencing. Nature biotechnology 26, 1135–1145 (2008).

23 Aebersold, R. & Mann, M. Mass spectrometry-based proteomics. Nature 422, 198–207 (2003).

24 Naumova, O. Y., Lee, M., Rychkov, S. Y., Vlasova, N. V. & Grigorenko, E. L. Gene expression in the human brain: the current state of the study of specificity and spatiotemporal dynamics. Child development 84, 76–88 (2013).

25 Kang, H. J. et al. Spatio-temporal transcriptome of the human brain. Nature 478, 483–489 (2011).

26 Lee, J. H. et al. Fluorescent in situ sequencing (FISSEQ) of RNA for gene expression profiling in intact cells and tissues. Nature protocols 10, 442–458 (2015).

27 Mortazavi, A., Williams, B. A., McCue, K., Schaeffer, L. & Wold, B. Mapping and quantifying mammalian transcriptomes by RNA-Seq. Nature methods 5, 621–628 (2008).

28 Roopenian, D. C. & Akilesh, S. FcRn: the neonatal Fc receptor comes of age. Nature reviews immunology 7, 715–725 (2007).

29 Rissin, D. M. et al. Single-molecule enzyme-linked immunosorbent assay detects serum proteins at subfemtomolar concentrations. Nature biotechnology 28, 595–599 (2010).

30 Swaminathan, J. et al. Highly parallel single-molecule identification of proteins in zeptomole-scale mixtures. Nature biotechnology 36, 1076–1082 (2018).

31 Farka, Z. et al. Advances in Optical Single-Molecule Detection: En Route to Supersensitive Bioaffinity Assays. Angewandte Chemie International Edition 59, 10746–10773 (2020).

32 Sze, J. Y., Ivanov, A. P., Cass, A. E. G. & Edel, J. B. Single molecule multiplexed nanopore protein screening in human serum using aptamer modified DNA carriers. Nature communications 8, 1552 (2017).

33 Tannous, B. A., Kim, D.-E., Fernandez, J. L., Weissleder, R. & Breakefield, X. O. Codon-optimized Gaussia luciferase cDNA for mammalian gene expression in culture and in vivo. Molecular Therapy 11, 435–443 (2005).

34 Aalipour, A. et al. Engineered immune cells as highly sensitive cancer diagnostics. Nature biotechnology 37, 531–539 (2019).

35 Deane, R. et al. IgG-assisted age-dependent clearance of Alzheimer’s amyloid β peptide by the blood–brain barrier neonatal Fc receptor. Journal of Neuroscience 25, 11495–11503 (2005).

36 Cooper, P. R. et al. Efflux of monoclonal antibodies from rat brain by neonatal Fc receptor, FcRn. Brain research 1534, 13–21 (2013).

37 Zhang, Y. & Pardridge, W. M. Mediated efflux of IgG molecules from brain to blood across the blood–brain barrier. Journal of neuroimmunology 114, 168–172 (2001).

38 Borrok, M. J. et al. pH-dependent binding engineering reveals an FcRn affinity threshold that governs IgG recycling. Journal of Biological Chemistry 290, 4282–4290 (2015).

39 Westerink, R. H. S. & Ewing, A. G. The PC12 cell as model for neurosecretion. Acta Physiologica 192, 273–285 (2008).

40 Verhaegen, M. & Christopoulos, T. K. Recombinant Gaussia luciferase. Overexpression, purification, and analytical application of a bioluminescent reporter for DNA hybridization. Analytical chemistry 74, 4378–4385 (2002).

41 Lee, C.-H. et al. An engineered human Fc domain that behaves like a pH-toggle switch for ultra-long circulation persistence. Nature communications 10, 5031 (2019).

42 Herculano-Houzel, S., Mota, B. & Lent, R. Cellular scaling rules for rodent brains. Proceedings of the National Academy of Sciences 103, 12138–12143 (2006).

43 Gil, G. A. et al. c-Fos activated phospholipid synthesis is required for neurite elongation in differentiating PC12 cells. Molecular biology of the cell 15, 1881–1894 (2004).

44 Alexander, G. M. et al. Remote control of neuronal activity in transgenic mice expressing evolved G protein-coupled receptors. Neuron 63, 27–39 (2009).

45 Roth, B. L. DREADDs for neuroscientists. Neuron 89, 683–694 (2016).

46 Szablowski, J. O., Lee-Gosselin, A., Lue, B., Malounda, D. & Shapiro, M. G. Acoustically targeted chemogenetics for the non-invasive control of neural circuits. Nature biomedical engineering 2, 475–484 (2018).

47 Sørensen, A. T. et al. A robust activity marking system for exploring active neuronal ensembles. elife 5, e13918 (2016).

48 Bernau, K. et al. In vivo tracking of human neural progenitor cells in the rat brain using bioluminescence imaging. Journal of neuroscience methods 228, 67–78 (2014).

49 Chan, K. Y. et al. Engineered AAVs for efficient noninvasive gene delivery to the central and peripheral nervous systems. Nature neuroscience 20, 1172–1179 (2017).

50 Zetterberg, H. & Burnham, S. C. Blood-based molecular biomarkers for Alzheimer’s disease. Molecular brain 12, 26 (2019).

51 Niederkofler, V. et al. Identification of serotonergic neuronal modules that affect aggressive behavior. Cell reports 17, 1934–1949 (2016).

52 Clifton, N. E. et al. Dynamic expression of genes associated with schizophrenia and bipolar disorder across development. Translational psychiatry 9, 74 (2019).

53 Tiklová, K. et al. Disease Duration Influences Gene Expression in Neuromelanin-Positive Cells From Parkinson’s Disease Patients. Frontiers in molecular neuroscience 14, 763777 (2021).

54 Berchtold, N. C. et al. Gene expression changes in the course of normal brain aging are sexually dimorphic. Proceedings of the National Academy of Sciences 105, 15605–15610 (2008).

55 Ham, S. & Lee, S.-J. V. Advances in transcriptome analysis of human brain aging. Experimental & Molecular Medicine 52, 1787–1797 (2020).

56 Pasi, K. J. et al. Multiyear follow-up of AAV5-hFVIII-SQ gene therapy for hemophilia A. The New England Journal of Medicine 382, 29–40 (2020).

57 Hampel, H. et al. Blood-based biomarkers for Alzheimer disease: mapping the road to the clinic. Nature Reviews Neurology 14, 639–652 (2018).

58 Nakamura, A. et al. High performance plasma amyloid-β biomarkers for Alzheimer’s disease. Nature 554, 249–254 (2018).

59 Iturria-Medina, Y., Khan, A. F., Adewale, Q., Shirazi, A. H. & Initiative, A. s. D. N. Blood and brain gene expression trajectories mirror neuropathology and clinical deterioration in neurodegeneration. Brain 143, 661–673 (2020).

60 Mondello, S. et al. Blood-based protein biomarkers for the management of traumatic brain injuries in adults presenting to emergency departments with mild brain injury: a living systematic review and meta-analysis. Journal of neurotrauma 38, 1086–1106 (2021).

61 Kingsmore, S. F. Multiplexed protein measurement: technologies and applications of protein and antibody arrays. Nature reviews Drug discovery 5, 310–321 (2006).

62 Sun, X. et al. Functionally distinct neuronal ensembles within the memory engram. Cell 181, 410-423.e417 (2020).

63 Alfaro, J. A. et al. The emerging landscape of single-molecule protein sequencing technologies. Nature methods 18, 604–617 (2021).

64 Ying, T., Feng, Y., Wang, Y., Chen, W. & Dimitrov, D. S. Monomeric IgG1 Fc molecules displaying unique Fc receptor interactions that are exploitable to treat inflammation-mediated diseases. MAbs 6, 1201–1210 (2014).

65 Challis, R. C. et al. Systemic AAV vectors for widespread and targeted gene delivery in rodents. Nature protocols 14, 379–414 (2019).

66 Lawlor, P. A., Bland, R. J., Mouravlev, A., Young, D. & During, M. J. Efficient gene delivery and selective transduction of glial cells in the mammalian brain by AAV serotypes isolated from nonhuman primates. Molecular therapy 17, 1692–1702 (2009).

