## Supplementary Information for "Engineered Serum Markers for Noninvasive Monitoring of Gene Expression in the Brain"

---

### List of Supplementary Materials

Supplementary Figure 1. RMA secretion assay. p. 2

Supplementary Figure 2. Additional brain images of mice intracranially injected with RMA proteins. p. 3

Supplementary Figure 3. Half-life of Gluc-RMA. p. 3

Supplementary Figure 4. Correlation of brain transduction efficiency to plasma Gluc-RMA signal. p. 4

Supplementary Figure 5. Detecting the *Fos* promoter activity in PC-12. p. 5

Supplementary Figure 6. Gluc-RMA detects neuronal activity of the caudate putamen. p. 6

Supplementary Figure 7. Gluc-RMA detects neuronal activity of the hippocampus. p. 7

Supplementary Figure 8. Expression of Gluc-RMA in the brain and periphery organs from i.v. PHP.eB delivery. p. 8

Supplementary Figure 9. Analysis of GFP positive cells using FACS. p. 9

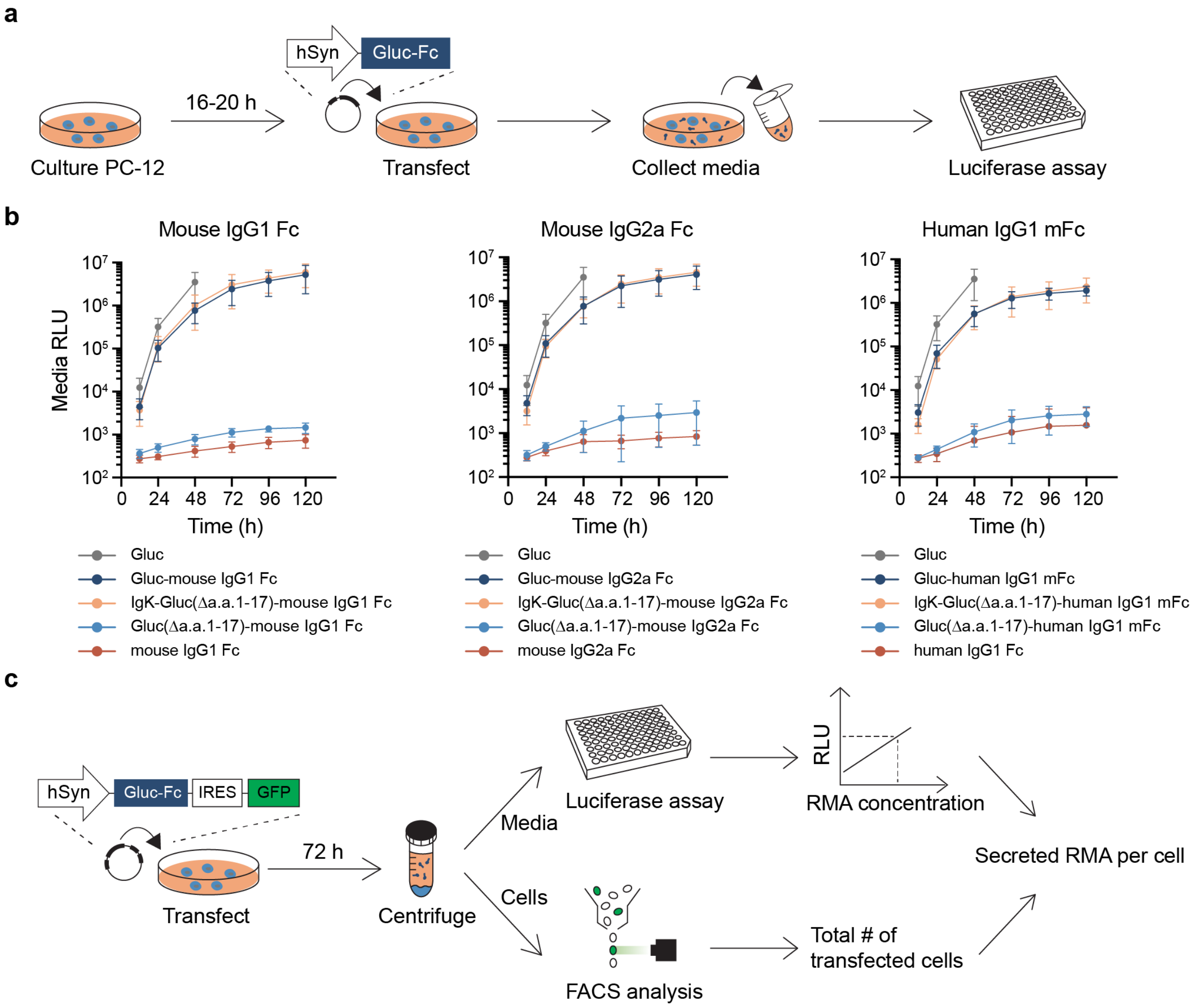


**Supplementary Figure 1 | RMA secretion assay. a,** Experimental scheme for detecting secreted RMA using PC-12. **b**, Signal-peptide dependency of RMA secretion. RMAs lacking the signal peptide are truncated by the first 17 amino acids of Gluc (Δa.a.1-17). Replacement of the native signal-peptide of Gluc with the murine Ig Kappa (Igκ) did not alter the secretion efficiency. *n*=5 independent cultures analyzed. Data are shown as mean ± SD. **c**, Workflow for estimating the number of RMA secreted per PC-12 cell. Delivering bicistronic vector co-expressing Gluc-Fc and GFP allows for measurement of the secreted RMAs and the number of transfected cells to calculate the average amount of RMA proteins released per cell. IRES: Internal ribosome entry site; FACS: Fluorescence-activated cell sorting.

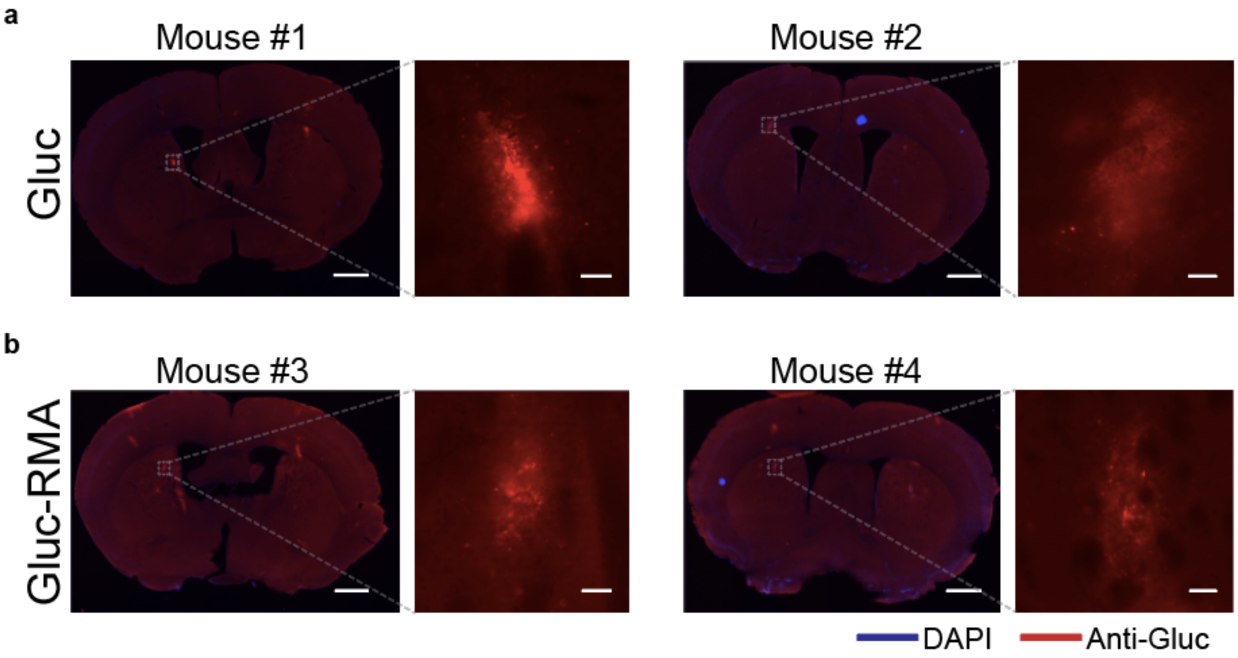


**Supplementary Figure 2 | Additional brain images of mice intracranially injected with RMA proteins. a**, and **b**, Images of the brain 24 hr after the injection of **a)** Gluc or **b)** Gluc-RMA proteins. Each image is obtained from an independent mouse. Scale, 1000 µm (whole-brain image), and 50 µm (enlarged image).

**a**

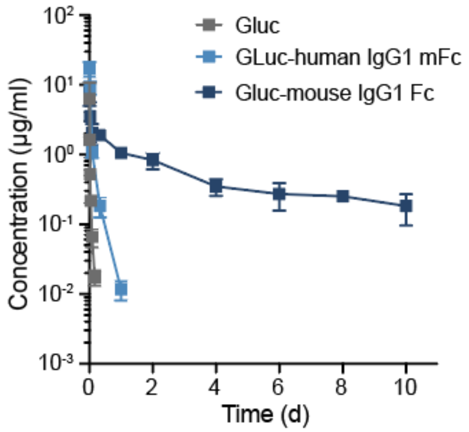


| Construct (protein) | $AUC_{\infty}$ (µg min ml$^{-1}$) | Clearance (ml min$^{-1}$ kg$^{-1}$) | α-phase $T_{1/2}$ (min) | β-phase $T_{1/2}$ (min) |
|---|---|---|---|---|
| Gluc | 127.3 ± 42.6 | 16.8 ± 4.7 | 7.2 ± 0.9 | 30.7 ± 2.9 |
| Gluc-human IgG1 Fc | 1224 ± 150 | 1.7 ± 0.2 | 29.1 ± 3.3 | 207.6 ± 19.0 |
| Gluc-mouse IgG1 Fc | 9685 ± 705 | 0.21 ± 0.01 | 120.9 ± 64.7 | 6269 ± 2283 |

**Supplementary Figure 3 | Half-life of Gluc-RMA. a,** Concentrations of Gluc, Gluc-human IgG1 mFc, and Gluc-mouse IgG1 Fc (Gluc-RMA) in the wild-type mice as a function of time after intravenous (i.v.) administration of each protein with the dose of 2 mg/kg. Two-compartment elimination model was used to analyze the pharmacokinetic parameters. *n*=2 to 6 blood samples analyzed. Data are shown as mean ± SD. AUC: Area under the curve.

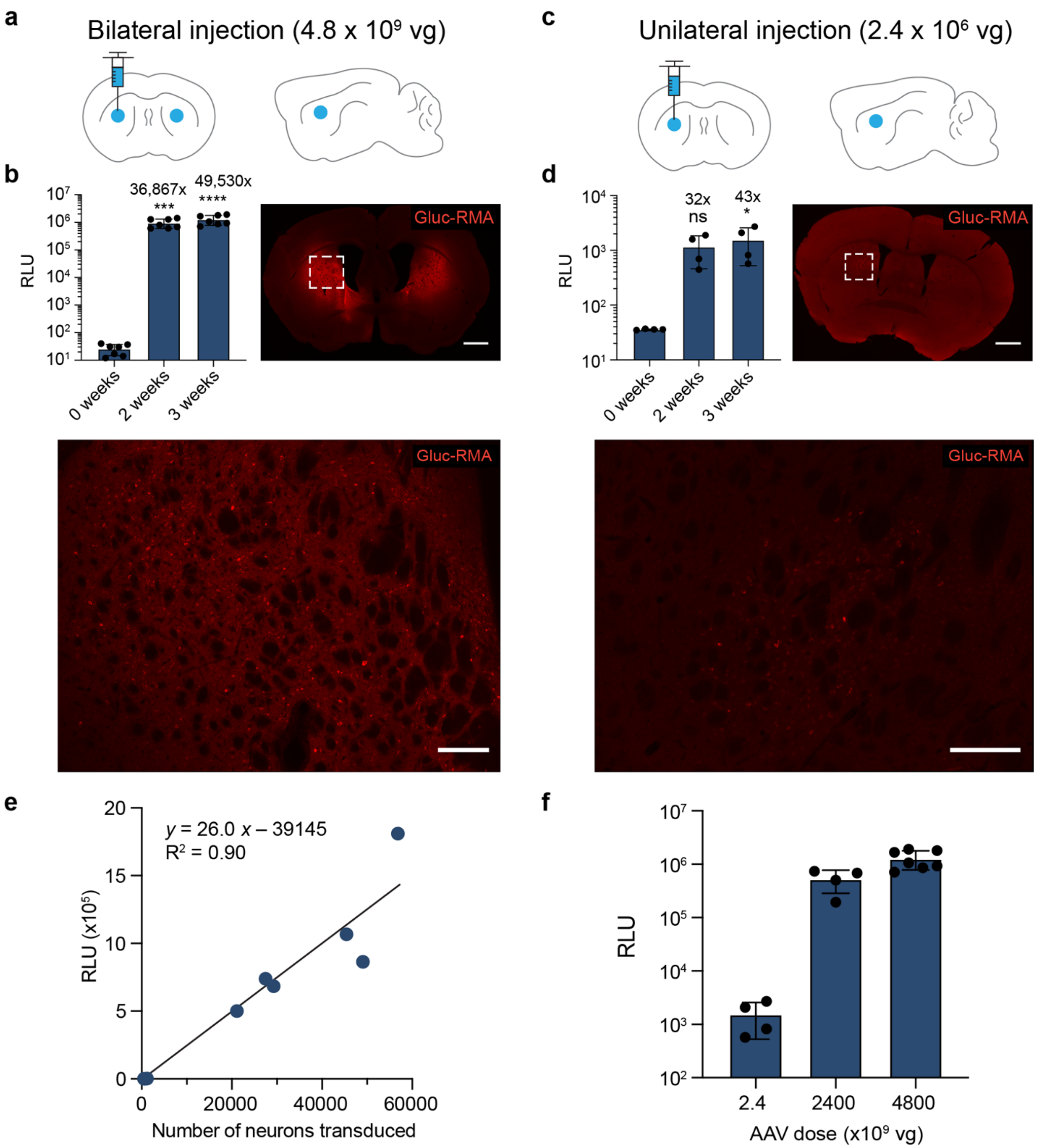


**Supplementary Figure 4 | Correlation of brain transduction efficiency to plasma Gluc-RMA signal. a** and **c**, Injection sites for the **a**) bilateral or **c**) unilateral delivery of AAV encoding Gluc-RMA into CP. The brain schemes display the coronal (left) and sagittal (right) views of the injection sites (blue circles). The AAV dose indicates total viral genomes injected per mouse. **b** and **d**, Plasma bioluminescence signal and representative brain images showing the Gluc-RMA gene expression. Left (bar graph): RLU measured from the collected blood samples. The number above each bar indicates the signal fold increase compared with the signal at 0 weeks. Right and bottom (Images): Brain slices stained against Gluc-RMA after 3 weeks post-AAV injection. Scale, 1000 µm (whole-brain image) and 200 µm (enlarged image). *$P$<0.05,

***$P$<0.001, ****$P$<0.0001, ns (not significant), in comparison with the plasma signal at 0 weeks, using one-way ANOVA, Tukey's test. Data are shown as mean ± SD. **e** and **f**, Correlation between the **e)** estimated number of transduced neurons, or **f)** AAV dose to the plasma bioluminescence signals.

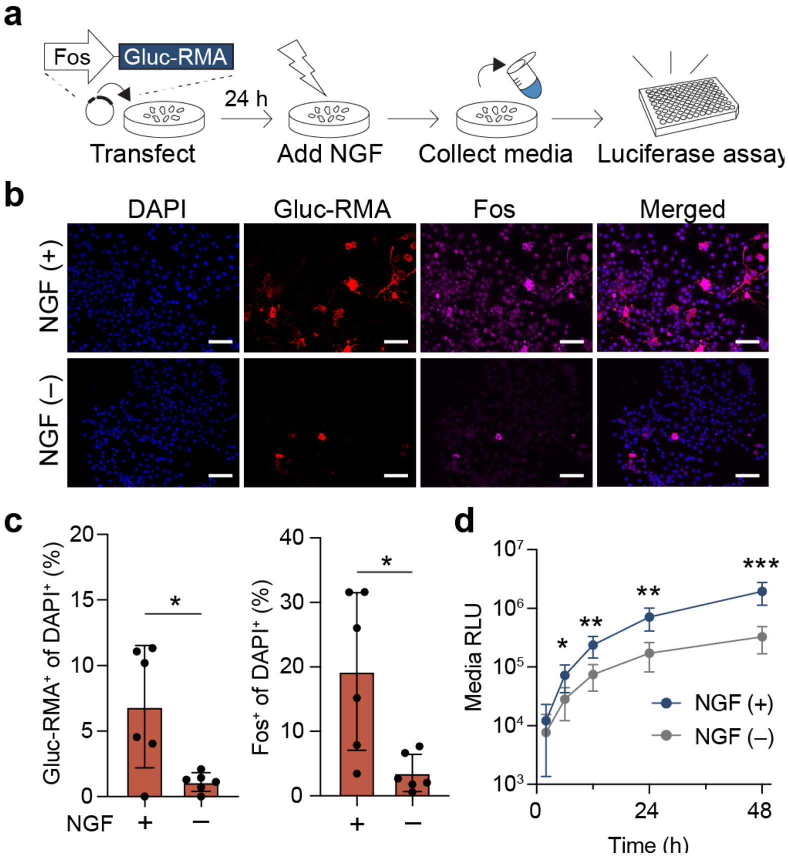


**Supplementary Figure 5 | Detecting the *Fos* promoter activity in PC-12. a**, Experimental scheme. PC-12 cells are transfected with plasmids encoding Gluc-RMA controlled under the *Fos* promoter, stimulated by NGF, and analyzed by luciferase assay for the secreted Gluc-RMA. **b**, Representative images of PC-12 stained 48 hr after the addition of media with or without NGF. Scale, 50 µm. **c**, Percentage of PC-12 cells that express Gluc-RMA (left) or *Fos* (right). **d**, RLU measured from the culture media that contains the released Gluc-RMA. Asterisks show the comparison between the RLU values with NGF (+) and without NGF (–) of the same time point. For **c** and **d**, $n$=6 independent cultures analyzed. *$P$<0.05, **$P$<0.01, ***$P$<0.001, using unpaired two-tailed t-test. Data are shown as mean ± SD. NGF: neural growth factor.

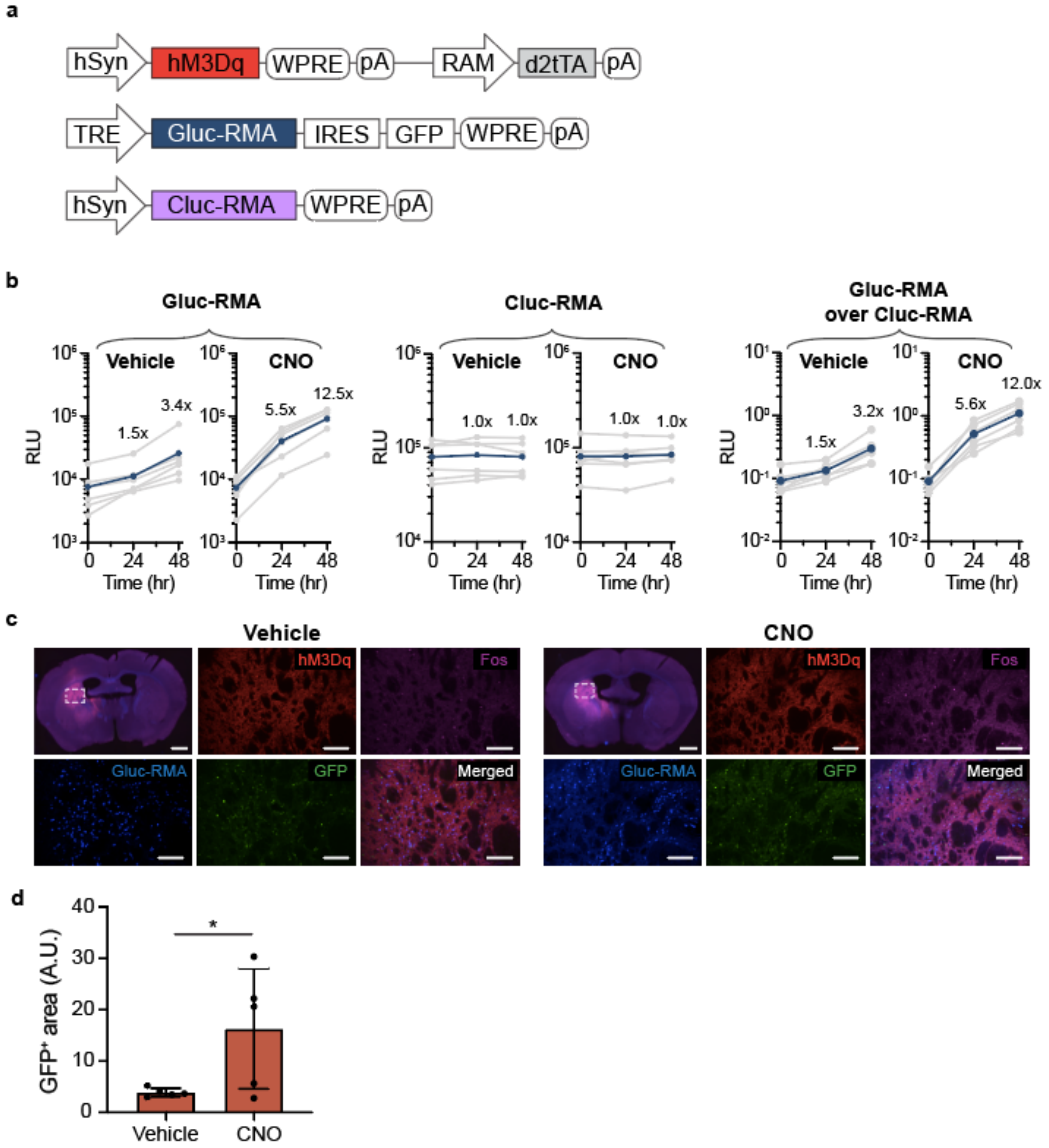


**Supplementary Figure 6 | Gluc-RMA detects neuronal activity of the caudate putamen. a**, AAV vector designs for the activatory DREADD, RAM-inducible Gluc-RMA, and Cluc-RMA expressed constitutively in neurons as a multiplexed signal and internal control. **b**, Average (blue) and individual mice (grey) plasma RLU values measured after the i.p. injection of either vehicle or 5 mg $kg^{-1}$ of CNO. Normalized signal (right) was calculated by dividing the RLU values of Gluc-RMA by that of Cluc-RMA of the same mouse at the same time point. *n*=6 independent mice analyzed. **c,** Representative images of stained brain slices of mouse perfused after 48 hr post-injection of either vehicle or CNO. Scale, 1000 μm (whole-brain image), 200 μm (enlarged image). **d**, Quantification of positive expression areas of GFP for images in Fig. 5d analyzed by the ImageJ software. *n*=5 independent samples analyzed. *$P$<0.05, in comparison between the vehicle- and CNO-injected groups, using unpaired two-tailed t-test. Data are shown as mean ± SD.

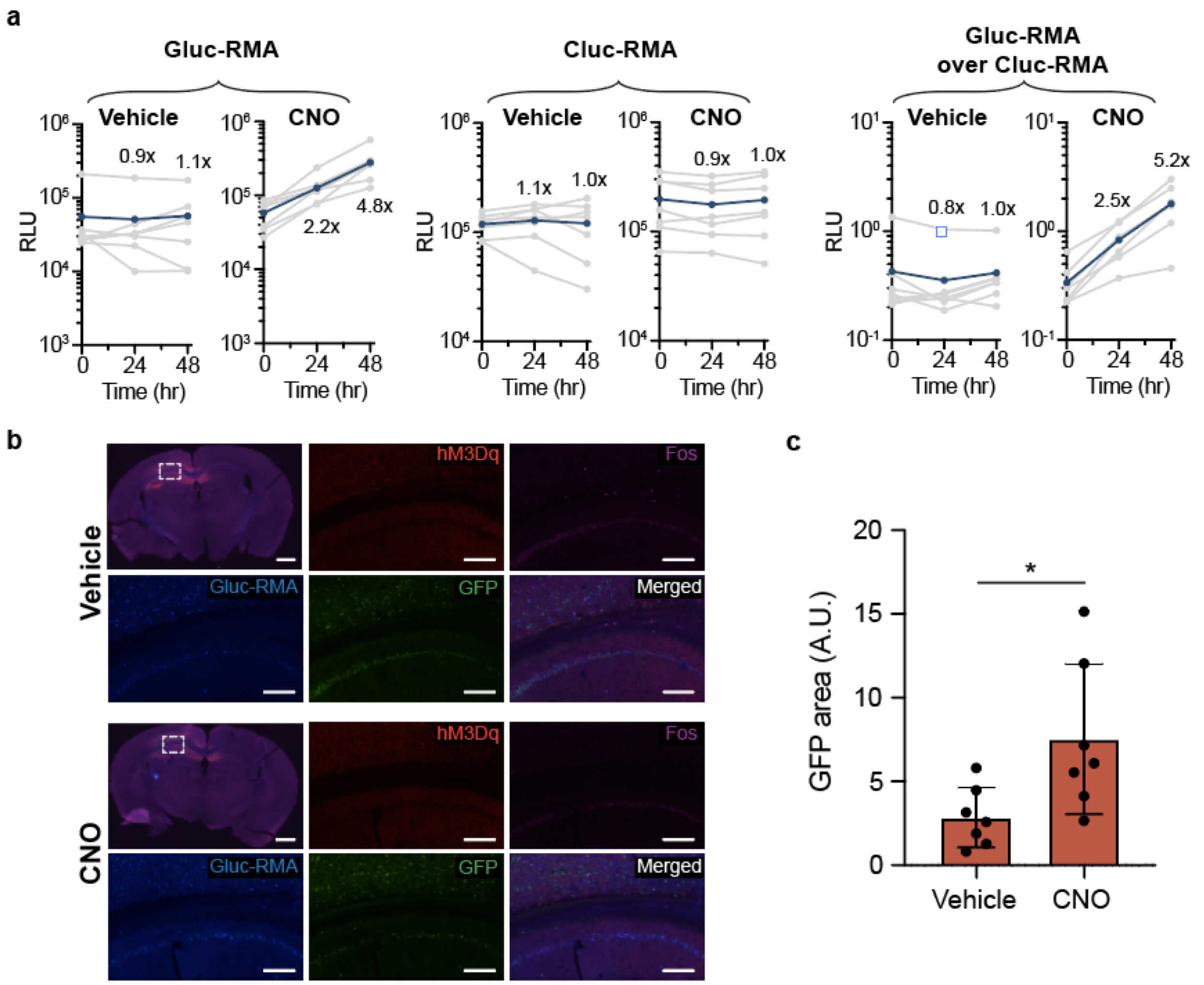


**Supplementary Figure 7 | Gluc-RMA detects neuronal activity of the hippocampus. a**, Average plasma RLU values (blue) calculated by those of individual mouse (light grey) measured after the i.p. injection of either vehicle or 5 mg $kg^{-1}$ of CNO. Normalized signal (right) was calculated by dividing the RLU value of Gluc-RMA by that of Cluc-RMA of the same mouse at the same time point. $n$=7 independent mice analyzed. **b**, Representative images of stained brain slices of mouse perfused after 48 hr post-injection. Scale, 1000 µm (whole-brain image), 200 µm (enlarged image). **c**, Quantification of positive expression areas of GFP analyzed by the ImageJ software. $n$=7 independent samples analyzed. *$P$<0.05, in comparison between the vehicle- and CNO-injected groups, using unpaired two-tailed t-test. Data are shown as mean ± SD.

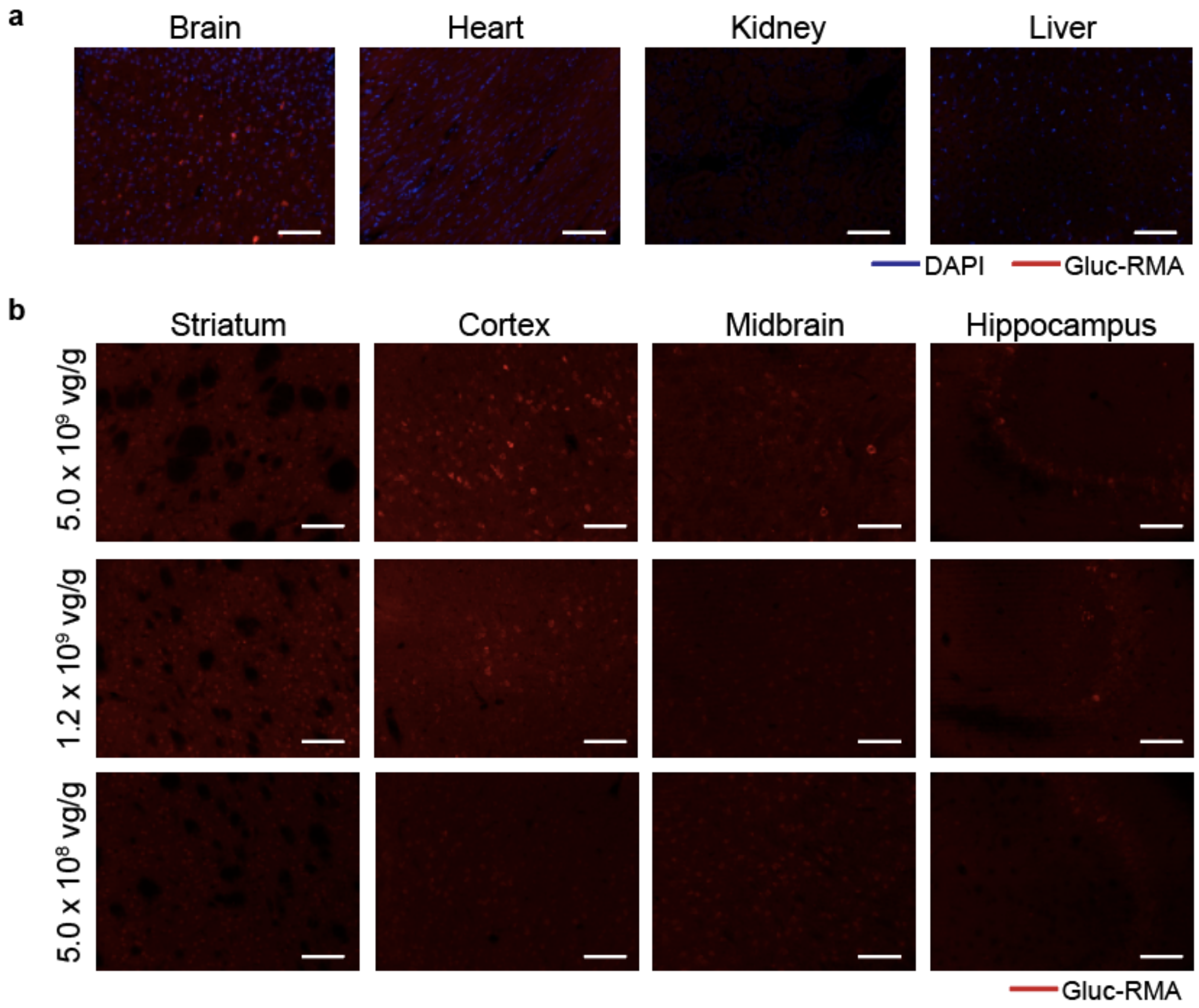


**Supplementary Figure 8 | Expression of Gluc-RMA in the brain and periphery organs from i.v. PHP.eB delivery. a**, Gluc-RMA expression in the brain, heart, kidney, and liver. Stained images show the expression occurs in the brain but not in other organs. Scale, 100 µm. **b**, Gluc-RMA expression in the striatum, cortex, midbrain, and hippocampus in the brain of mice administered with different PHP.eB doses. Scale, 100 µm.

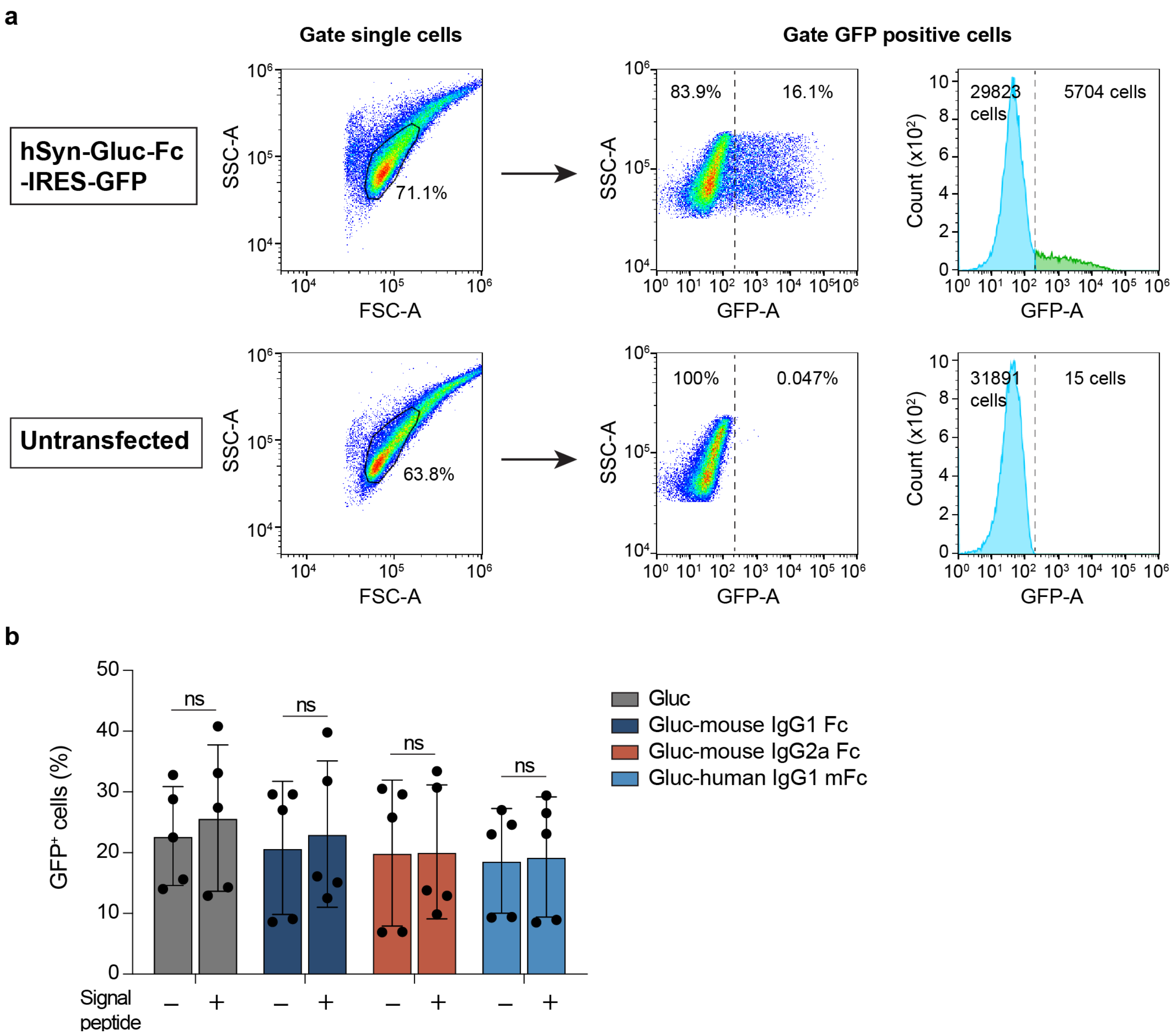


**Supplementary Figure 9 | Analysis of GFP positive cells using FACS. a**, Gating strategy for estimating the percentage of GFP positive cells. PC-12 cells were transfected with bicistronic vector encoding RMA and GFP. After 72 h post-delivery, spectral cell analyzer was used to sequentially gate single cells and then the GFP positive cells against the untransfected control. Above data represent a sample of PC-12 cells expressing Gluc-mouse IgG1 Fc and GFP versus the untransfected control. **b**, Percentage of PC-12 cells expressing GFP analyzed by FACS. ns (not significant) in comparison withe the percent GFP positive cells of the respective RMA, using two-way ANOVA Sidak's test. Data are shown as mean ± SD.
